# Individuals’ expected genetic contributions to future generations, reproductive value, and short-term metrics of fitness in free-living song sparrows (Melospiza melodia)

**DOI:** 10.1101/528513

**Authors:** Jane M. Reid, Pirmin Nietlisbach, Matthew E. Wolak, Lukas F. Keller, Peter Arcese

## Abstract

Appropriately defining and enumerating ‘fitness’ is fundamental to explaining and predicting evolutionary dynamics. Yet theoretical concepts of fitness are often hard to translate into quantities that can be quantified in wild populations experiencing complex environmental, demographic, genetic and selective variation. While the ‘fittest’ entities might be widely understood to be those that ultimately leave most descendants at some future time, such long-term legacies are hard to measure, impeding evaluation of how well more tractable short-term metrics of individual fitness directly predict longer-term outcomes. One opportunity for conceptual and empirical convergence stems from the principle of individual reproductive value (*V*_i_), defined as the number of copies of each of an individual’s alleles that is expected to be present in future generations given the individual’s realised pedigree of descendants. Since *V*_i_ tightly predicts an individual’s longer-term genetic contribution, quantifying *V*_i_ provides a tractable route to quantifying what, to date, has been an abstract fitness concept. We used complete pedigree data from free-living song sparrows (*Melospiza melodia*) to demonstrate that individuals’ expected genetic contributions stabilise within an observed 20-year time period, allowing individual *V*_i_ to be evaluated. Considerable among-individual variation in *V*_i_ was evident in both sexes. However, standard short-term metrics of individual fitness, comprising lifespan, lifetime reproductive success and projected growth rate, typically explained less than half the variation. Given these results, we discuss what evolutionary inferences can and cannot be directly drawn from short-term versus longer-term fitness metrics observed on individuals, and highlight that analyses of pedigree structure may provide useful complementary insights into evolutionary processes and outcomes.

## Introduction

Appropriately defining and enumerating ‘fitness’ is fundamental to all theoretical and empirical attempts to explain and predict the dynamics of allele frequencies, phenotypes and populations (de Jong 1994; Day & Otto 2001; Grafen 2006; Orr 2009; Sæther & Engen 2015). Yet it has proved hard to define quantitative metrics of fitness that unify all theoretical and empirical sub-disciplines in evolutionary biology, and to translate theoretically useful concepts into quantities that can feasibly be measured in wild populations (Kozłowski 1993; de Jong 1994; Käär & Jokela 1998; Brommer 2000; Metcalf & Parvard 2007; Orr 2009; Hunt & Hodgson 2010; Sæther & Engen 2015; Grafen 2018). Perhaps the closest to overarching conceptual unanimity is the broad idea that the ‘fittest’ alleles, genotypes or individuals are ultimately those that contribute most descendants to a population at some point in the future (Benton & Grant 2000; Day & Otto 2001; Brommer *et al.* 2002, 2004; Hunt *et al.* 2004; Grafen 2006; Roff 2008; Hunt & Hodgson 2010; Graves & Weinreich 2017). Yet, such concepts can seem remote from the short-term metrics of individual fitness that empiricists working on free-living populations commonly aim to measure, which typically comprise simple functions of individuals’ realised survival and/or reproductive success (Brommer *et al.* 2002, 2004; Link *et al.* 2002; Coulson *et al.* 2006; Hendry *et al.* 2018; Wolak *et al.* 2018). Such metrics can correctly enumerate individual contributions to the next year or generation, but will not necessarily directly predict longer-term genetic contributions, especially given compound density-, frequency- and/or environment-dependent selection (de Jong 1994; Day & Otto 2001; Hunt *et al.* 2004; Roff 2008; Sæther & Engen 2015; Graves & Weinreich 2017).

One opportunity for conceptual and empirical convergence stems from the principle of individual ‘reproductive value’ (*V*_i_), defined here as the number of copies of each of an individual’s alleles that is expected to be present in future generations given the individual’s realised pedigree of descendants (Barton & Etheridge 2011). *V*_i_ results from Mendelian allele inheritance given the realised survival and reproductive success of a focal individual and successive generations of descendants, which in turn result from context-dependent natural and sexual selection on multivariate expressed phenotypes alongside environmental and demographic stochasticity. In sexually-reproducing species, any successful individual’s ultimate genetic contribution will emerge over long timeframes (i.e. >>100 generations), but its expected genetic contribution (i.e. *V*_i_) should stabilise over relatively few generations: approximately log_2_(*N*) where *N* is population size (Chang 1999; Barton & Etheridge 2011). This equals approximately 7, 10 and 13 generations given *N* = 100, 1000 and 10000 respectively, meaning that *V*_i_ should be estimable within timeframes that are increasingly within reach of empirical studies of free-living populations, at least for species with moderately short generation times. Further, Barton & Etheridge (2011) showed that an individual’s stabilised *V*_i_ accurately predicts the longer-term probability that a neutral or weakly selected allele carried by that individual will persist in the focal population (versus go extinct). Importantly, this holds true even at the level of single individuals and alleles, not just groups. Consequently, an individual’s stabilised *V*_i_ tightly predicts its longer-term genetic contribution (Barton & Etheridge 2011).

These key theoretical results were initially derived assuming idealised conditions of neutral alleles segregating in well-mixed (i.e. random mating, no population structure or subdivision) diploid Wright-Fisher populations with discrete non-overlapping generations and constant population sizes that are sufficiently large to preclude short-term inbreeding. However, they also apply adequately given some population structure and different distributions of reproductive success, and to alleles that affect reproductive success (given standard assumptions of the infinitesimal model, implying weak selection, Chang 1999; Barton & Etheridge 2011; Barton *et al.* 2017). To the degree that such short-term stabilisation and associated predictive ability of *V*_i_ hold under natural conditions encompassing age-structure, overlapping generations, non-random mating, highly heterogeneous reproductive success within and between lineages, and dynamic finite population sizes with inbreeding (Chang 1999; Gravel & Steel 2015), then the availability of multi-generation pedigree data from wild populations provides opportunities to directly quantify *V*_i_ and thereby infer longer-term probabilities of allele persistence and individual genetic contributions. This provides a route to direct field estimation of what has, to date, been an abstract theoretical fitness concept.

Datasets that allow estimation of *V*_i_ can then be used to examine the degree to which more typically tractable short-term metrics of individual fitness, comprising simple functions of individuals’ realised survival and reproductive success can predict *V*_i_ and hence longer-term individual genetic contributions, and thereby evaluate what evolutionary inferences can or cannot be directly drawn from such metrics (Brommer *et al.* 2002, 2004). Widely-used metrics of individual fitness include lifetime reproductive success (LRS), a time-independent metric defined as the total number of offspring produced by an individual over its lifetime, and individual growth rate (λ_ind_), a time-dependent metric that emphasises offspring produced early in life (McGraw & Caswell 1996; Käär & Jokela 1998; Brommer *et al.* 2002, 2004; MacColl & Hatchwell 2004, see Methods). In principle, LRS and λ_ind_ should equally well predict individual long-term genetic contributions given constant population size and lineage-invariant selection, but have different expected utilities otherwise (λ_ind_ may be more appropriate in increasing populations, Brommer *et al.* 2002; Graves & Weinreich 2017). Both metrics are expected to out-perform measures of individual survival or lifespan, especially given trade-offs between survival and reproduction (Day & Otto 2001; Brommer *et al.* 2002; Hunt *et al.* 2004).

Yet, in reality, populations do not retain constant growth rates, but vary in size due to environmental stochasticity, and experience density-, frequency- and/or environment-dependent selection alongside substantial within-lineage demographic stochasticity (de Jong 1994; Benton & Grant 2000; Hunt *et al.* 2004; Lande *et al.* 2009). Allele frequency dynamics then become very difficult to predict, especially in age-structured populations, even if key rules governing selection and population dynamics are known (Day & Otto 2001; Lande *et al.* 2009; Gravel & Steel 2015; Sæther & Engen 2015; Myhre *et al.* 2016; Graves & Weinreich 2017). The degree to which short-term metrics of individual fitness do (or do not) predict *V*_i_ and hence longer-term individual genetic contributions given natural demographic variation, and thereby allow direct long-term evolutionary inference, then becomes an empirical question (Brommer *et al.* 2002, 2004; Hunt & Hodgson 2010; Graves & Weinreich 2017).

We use >20 years of complete genetically-verified pedigree data from free-living song sparrows (*Melospiza melodia*) to evaluate the degree to which individuals’ expected genetic contributions stabilise within the observed number of years and generations, thereby allowing individual *V*_i_ to be estimated and longer-term probabilities of allele persistence and genetic contributions to be inferred. We then quantify the degree to which individuals’ stabilised *V*_i_ values are predicted by standard short-term metrics of individual fitness (lifespan, LRS and λ_ind_), and thereby elucidate what can be directly inferred from such metrics in the context of natural environmental, demographic, genetic and selective variation.

## Methods

### Study system

Quantifying individual *V*_i_ (the expected number of allele copies contributed to future generations conditional on an individual’s realised pedigree of descendants, Barton & Etheridge 2011) requires complete, accurate, pedigree data spanning sufficient generations (approximately log_2_(*N*)) for the expectation to stabilise. This equates to approximately *G*log_2_(*N*) years, where *G* is mean generation time. Such data exist for a small, resident, population of song sparrows inhabiting Mandarte island, British Columbia, Canada (Supporting Information S1). The available data allow calculation of any desired metric of relatedness and fitness across individuals hatched from 1992, with most likely zero error or missing data with respect to the local population (Reid *et al.* 2014, 2016; Wolak *et al.* 2018).

Among-year variation in local environmental conditions and population density drives considerable among-year variation in song sparrow reproduction and survival (Arcese *et al.* 1992; Wilson & Arcese 2003; Tarwater *et al.* 2018), inducing substantial among-cohort variation in mean lifespan and LRS (Lebigre *et al.* 2012; Wolak *et al.* 2018). Total adult population size has consequently varied substantially among years (arithmetic mean: 73±29SD individuals, range: 33-128, Supporting Information S1). The adult sex ratio was typically male-biased (mean proportion males: 0.59±0.07SD, range: 0.39-0.75, Supporting Information S1), allowing the mean and variance in LRS to differ between females and males (Lebigre *et al.* 2012). Generation time, calculated as mean parent age, is approximately 2.5 years (Supporting Information S2). Following basic theory (Chang 1999), individuals’ expected genetic contributions should therefore stabilise within roughly *G*log_2_(*N*) ≈ 2.5log_2_(73) ≈ 15 years, and potentially sooner since harmonic mean population size and effective population size (*N*_e_) are less than the arithmetic mean (O’Connor *et al.* 2006, Supporting Information S1). Stabilised *V*_i_ should therefore be estimable for song sparrows hatched early within the period for which genetically-verified pedigree data for descendants are currently available (i.e. 1992-2015).

### Calculation of individuals’ expected genetic contributions

The first objective was to calculate the number of copies of a (hypothetical) autosomal allele present in any focal individual that is expected to be present in the focal population in each year following the focal individual’s natal year, and thereby evaluate each individual’s stabilised *V*_i_. Such expectations can be calculated directly (i.e. analytically) from pedigree data using standard recursive algorithms for coefficients of kinship (Caballero & Toro 2000; Reid *et al.* 2016). However, the full distributions of allele copy numbers, and hence the variance around the expectation and associated probabilities of allele extinction, cannot be straightforwardly calculated for individuals in complex pedigrees with irregular inbreeding. We therefore used ‘gene-dropping’ simulations on the observed pedigree to compute key quantities (e.g. MacCluer *et al.* 1986; Caballero & Toro 2000).

Since song sparrows have considerably overlapping generations (median age at first reproduction: 1 year; maximum lifespan: 10 years, Supporting Information S2), analyses focussed on cohorts and years rather than discrete generations. Each individual hatched in a focal cohort that survived to adulthood (i.e. age one year) was assigned a unique allele identity, which was ‘dropped’ down the observed pedigree assuming autosomal Mendelian inheritance (i.e. passed to each offspring of each sex with probability 0.5). The identities of all alleles present in all individuals in the total extant population (i.e. all adults and ringed chicks, constituting a post-breeding census) in each subsequent year were extracted. Gene-drops were replicated 8,000 times. The mean number of copies of each allele present in each year (i.e. the expectation), the frequency of zero copies (and hence the probability of allele extinction, *P*_E_), and the mean, variance and coefficient of variance (CV=standard deviation/mean) of allele copy number conditional on allele persistence (i.e. number of copies >0) were computed across replicates. Allele copies present in descendants of both sexes were counted irrespective of the sex of the focal individual in which an allele originated, thereby tracking propagation of a hypothetical autosomal allele across the full pedigree rather than solely through single-sex lineages (contra Brommer *et al.* 2004). Current simulations therefore assume that focal cohort individuals are heterozygous and unrelated at the hypothetical locus (i.e. single unique alleles), and correctly account for realised within-lineage inbreeding in subsequent generations. They can therefore be considered to quantify the fate of a single neutral or weakly selected mutation in each focal individual (e.g. Barton & Etheridge 2011). Comparisons showed that gene-drop expectations diverged from analytical expectations by <1% on average. Gene-dropping was not applied to focal individuals that died before adulthood because their direct genetic contribution is known to be zero (hence *P*_E_=1), allowing them to be included in subsequent analyses without need for gene-drop computations.

### Analyses of individual genetic contributions and reproductive value

Gene-dropping focussed on song sparrow cohorts hatched in 1992-1994, for which ≥20 years of complete genetically-verified pedigree data on descendants are in hand (Supporting Information S1). To evaluate whether individual *V*_i_ could be reliably ascertained, we examined whether individuals’ absolute or proportional expected genetic contributions to the total extant population in each year stabilised within an observed 20-year post-hatch timeframe. Proportional contributions were calculated by dividing each individual’s absolute contribution (i.e. the expected number of allele copies) by the total number of alleles present in the population in the focal year (i.e. twice the total extant population size). This standardisation facilitates direct comparison of values across years and cohorts given varying population size. Stabilisation was evaluated by calculating the Pearson correlation coefficient (*r*_p_) between individuals’ gene-dropped expectations 20 years post-hatch versus in each previous year: convergence of *r*_p_ to one indicates stabilisation.

We then quantified the degree to which an individual’s expected genetic contribution 20 years post-hatch (interpreted as its *V*_i_) predicts key attributes of the full distribution of allele copy number emerging across gene-drop iterations. We examined the degree to which patterns concur with Barton & Etheridge’s (2011) theoretical development, supporting the premise that *V*_i_ predicts longer-term genetic contribution. First, to verify that *V*_i_ tightly predicts individual-level probability of allele extinction within the observed 20-year timeframe, and is consequently likely to do so across longer timeframes, we regressed the gene-dropped probability of allele extinction (*P*_E_) after 20 years on *V*_i_ across individuals. Second, to quantify the degree to which *V*_i_ also predicts the full distribution of allele copy number conditional on allele persistence, we related the mean, variance and CV of the number of copies after 20 years across gene-drop iterations where the allele did not go extinct to *V*_i_.

### Calculation of short-term metrics of individual fitness

The second objective was to evaluate the degree to which standard short-term metrics of individual realised fitness predict *V*_i_ and hence could in principle be used to directly infer individuals’ longer-term genetic contributions. In the context of evolutionary quantitative genetics, fitness metrics should ideally be measured zygote-to-zygote across a single generation (Lande & Arnold 1983; Wolf & Wade 2001; Hunt & Hodgson 2010). However, in practice, and in other contexts, they are commonly measured adult-to-offspring or adult-to-adult. This includes studies that aim to quantify fitness consequences of expression of adult traits (including reproductive or secondary sexual traits) and directly infer evolutionary outcomes, or to estimate *N*_e_ (e.g. Wolf & Wade 2001; Kokko *et al.* 2003; Hunt *et al.* 2004; MacColl & Hatchwell 2004; Sæther & Engen 2015; Myhre *et al.* 2016; Wolak *et al.* 2018). To encompass this spectrum, we extracted six fitness metrics for each focal individual. Lifespan was calculated as an individual’s age in its last observed summer (one metric per individual). LRS was calculated as the total numbers of ringed (i.e. 6 days post-hatch), independent (i.e. alive at cessation of parental care approximately 24 days post-hatch) or recruited (i.e. age one year) genetic offspring produced by each focal individual over its lifetime (hence three metrics of LRS per individual). λ_ind_ was calculated as the dominant eigenvalue of an individual projection matrix of dimension equal to the individual’s lifespan (McGraw & Caswell 1996; Brommer *et al.* 2002, Supporting Information S3). Top row fecundity terms were specified as either 0.5.m_ring_.ф_j_, where m_ring_ is the number of ringed offspring produced by each focal individual at each age and ф_j_ is the mean population-wide juvenile survival rate in the focal year, or as 0.5.m_rec_, where m_rec_ is the number of recruited offspring produced by each focal individual at each age (hence two metrics of λ_ind_ per individual, Supporting information S3). The factor 0.5 accounts for the transmission probability of a focal parental allele to each offspring given Mendelian inheritance. This must be directly incorporated into the calculation of λ_ind_, but can be readily applied as post-hoc scaling factor for analyses of LRS (Brommer *et al.* 2004; Reid *et al.* 2016).

### Analyses of individual fitness and reproductive value

To evaluate the degree to which the six short-term metrics of individual fitness explained and predicted variation in *V*_i_, we calculated Pearson and Spearman correlation coefficients, and linear regression slopes and associated adjusted R2 values, between *V*_i_ and each fitness metric across individuals. These statistics were calculated using individuals’ absolute fitness and *V*_i_, and using relative fitness and *V*_i_ (i.e. individual value divided by the sex-specific mean). Statistics were calculated across individuals from focal cohorts that survived to adulthood (mirroring common practice in studies that examine variation in fitness associated with adult phenotypes), and recalculated including values of zero for individuals that died before adulthood (thereby incorporating the otherwise ‘missing fraction’).

Because estimates of *V*_i_ for individuals hatched in 1992-1994 are partially non-independent (since pedigrees are partially nested, Supporting Information S1), and because we considered multiple non-independent fitness metrics and tests, we report estimated correlation and regression parameters but do not focus on statistical ‘significance’. Regression intercepts were estimated for analyses of lifespan but forced through the origin otherwise, because individuals with zero LRS or λ_ind_ have exactly zero *V*_i_. Since the total extant population sizes were similar across the three end years (i.e. 20 years post-hatch for each cohort, Supporting Information S1), analyses of absolute and proportional *V*_i_ yielded similar conclusions (Supporting Information S4 & S5). Key results were also quantitatively unchanged if gene-drops for all cohorts were run to the same end year (i.e. 22, 21 and 20 years post-hatch for the 1992, 1993 and 1994 cohorts respectively). Standard statistics (mean, variance, skew, CV) were used to summarise distributions of estimated *V*_i_ and fitness metrics. Analyses were implemented in R V3.3.3 (R Core Team 2017), using package nadiv (Wolak 2012).

## Results

### Individual genetic contributions and reproductive value

In total, 21, 24 and 10 female and 38, 23 and 23 male song sparrows hatched in 1992, 1993 and 1994 respectively survived to adulthood (i.e. age one year). These individuals’ absolute and proportional expected genetic contributions to the total extant population on Mandarte in each subsequent year clearly stabilised within the observed 20 year timeframe (Figs. 1 & 2, Supporting Information S4 & S5). Quantitatively, the correlations between the genetic contributions expected after 20 years and in each preceding year exceeded 0.95 by 12-13 years post-hatch in both sexes, and were typically close to one by the basic theoretical expectation of *∼*15 years post-hatch (Fig. 3). This implies that the expectations evident by 20 years post-hatch, and indeed from ∼13 years post-hatch, can be interpreted as good approximations of individual *V*_i_ (sensu Barton & Etheridge 2011).

**Figure 1.**
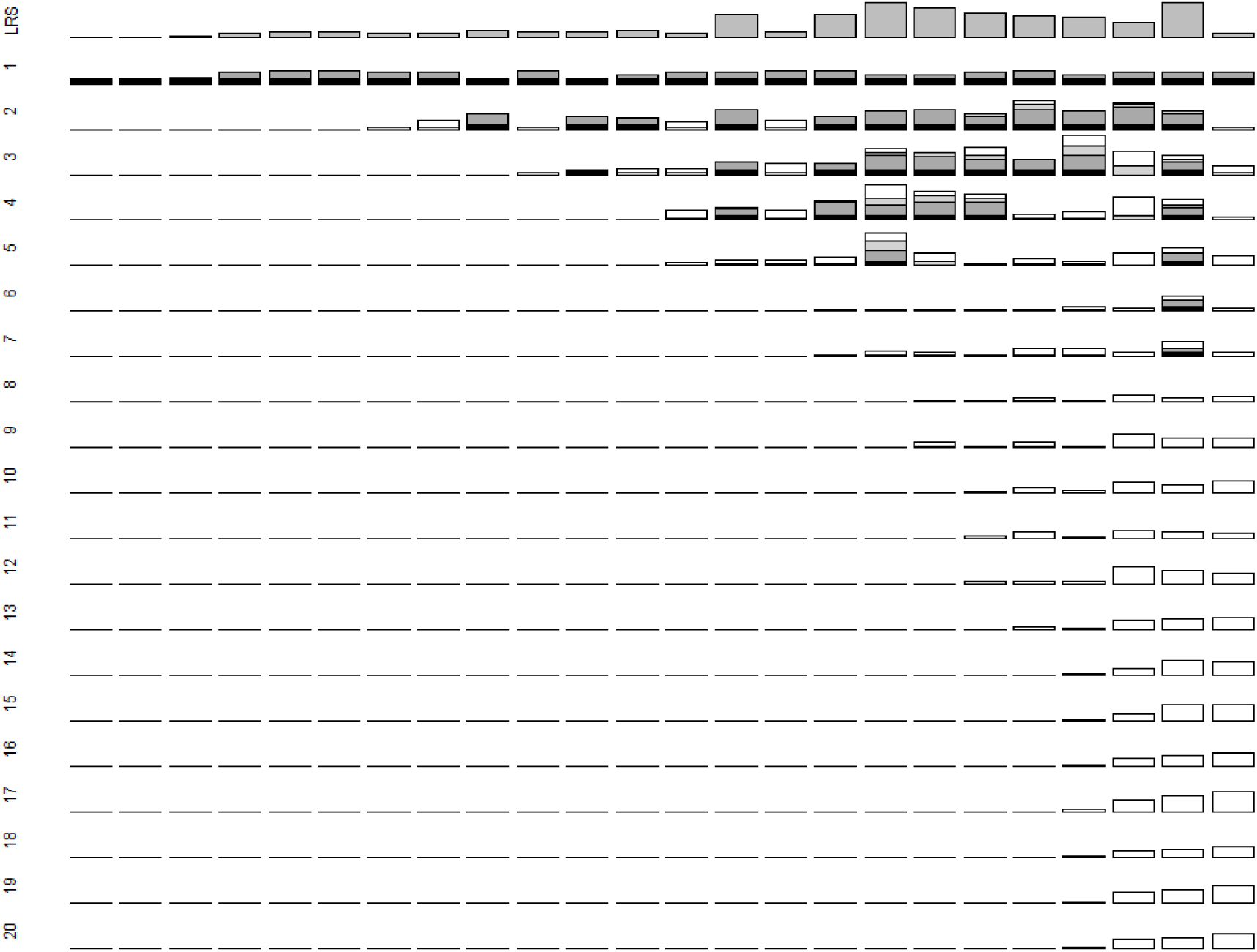
Observed lifetime reproductive success (LRS) measured as ringed offspring (top row) and absolute expected genetic contributions of 24 individual female song sparrows (columns) hatched in 1993 that survived to adulthood to the total extant population 1-20 years post-hatch (descending rows). Black shading denotes genetic contributions arising because a focal female was still alive in the focal year. Dark grey, light grey and white shading denote expected genetic contributions to offspring produced in the focal year, to surviving offspring produced in previous years, and to all subsequent descendants respectively. All bars (except LRS) are scaled to maximum y-axis values of 8 allele copies to allow direct comparison across years. Columns (i.e. females) are ordered by increasing expected contributions across final observed years. Equivalent data for females hatched in 1992 and 1994, and proportional genetic contributions for all females, are shown in Supporting Information S4 & S5.

**Figure 2.**
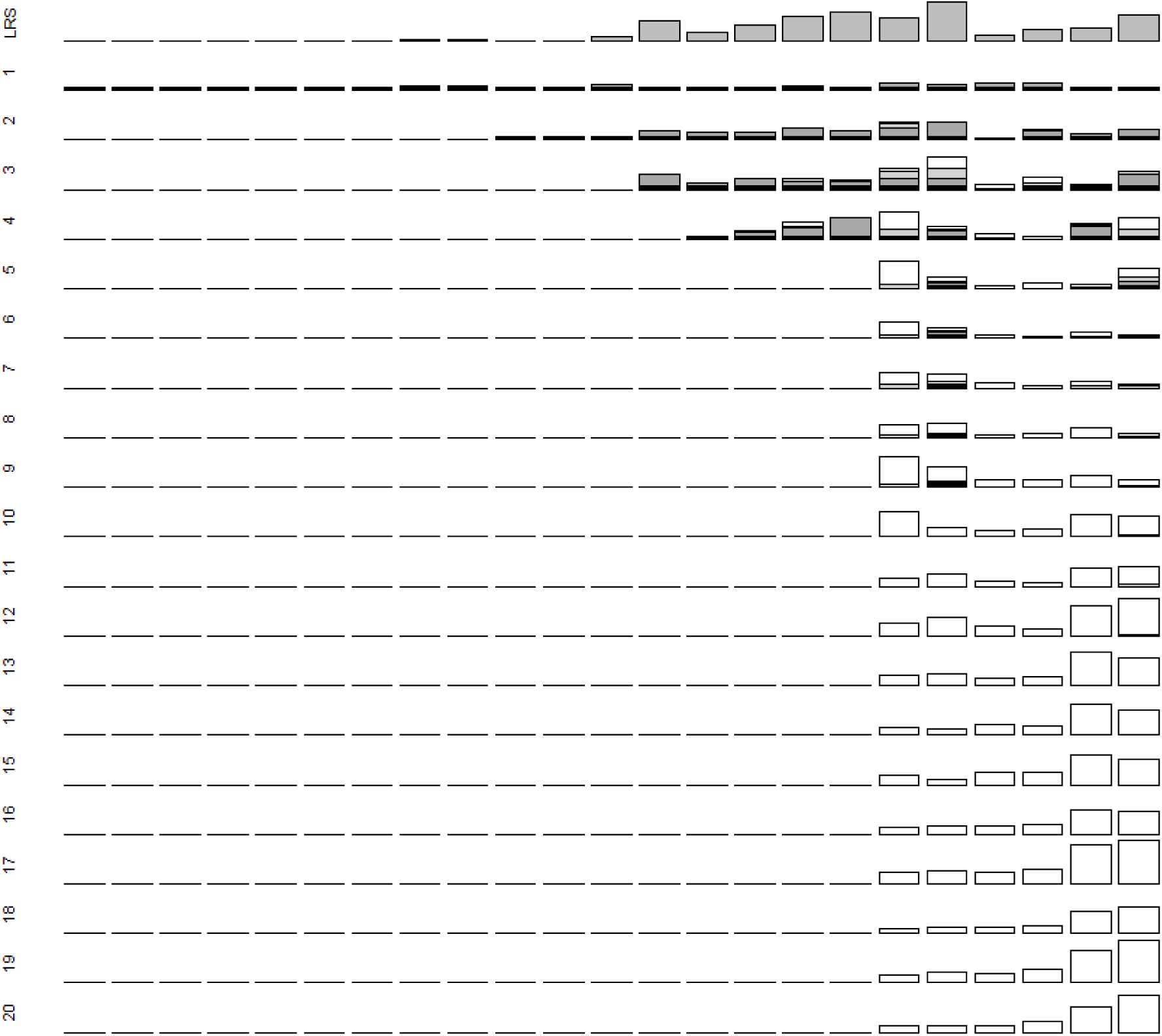
Observed lifetime reproductive success (LRS) measured as ringed offspring (top row) and absolute expected genetic contributions of 23 individual male song sparrows (columns) hatched in 1993 that survived to adulthood to the total extant population 1-20 years post-hatch (descending rows). Figure attributes are as for Figure 1. All bars (except LRS) are scaled to maximum y-axis values of 12 allele copies to allow direct comparison across years. Equivalent data for males hatched in 1992 and 1994, and proportional genetic contributions for all males, are shown in Supporting Information S4 & S5.

**Figure 3.**
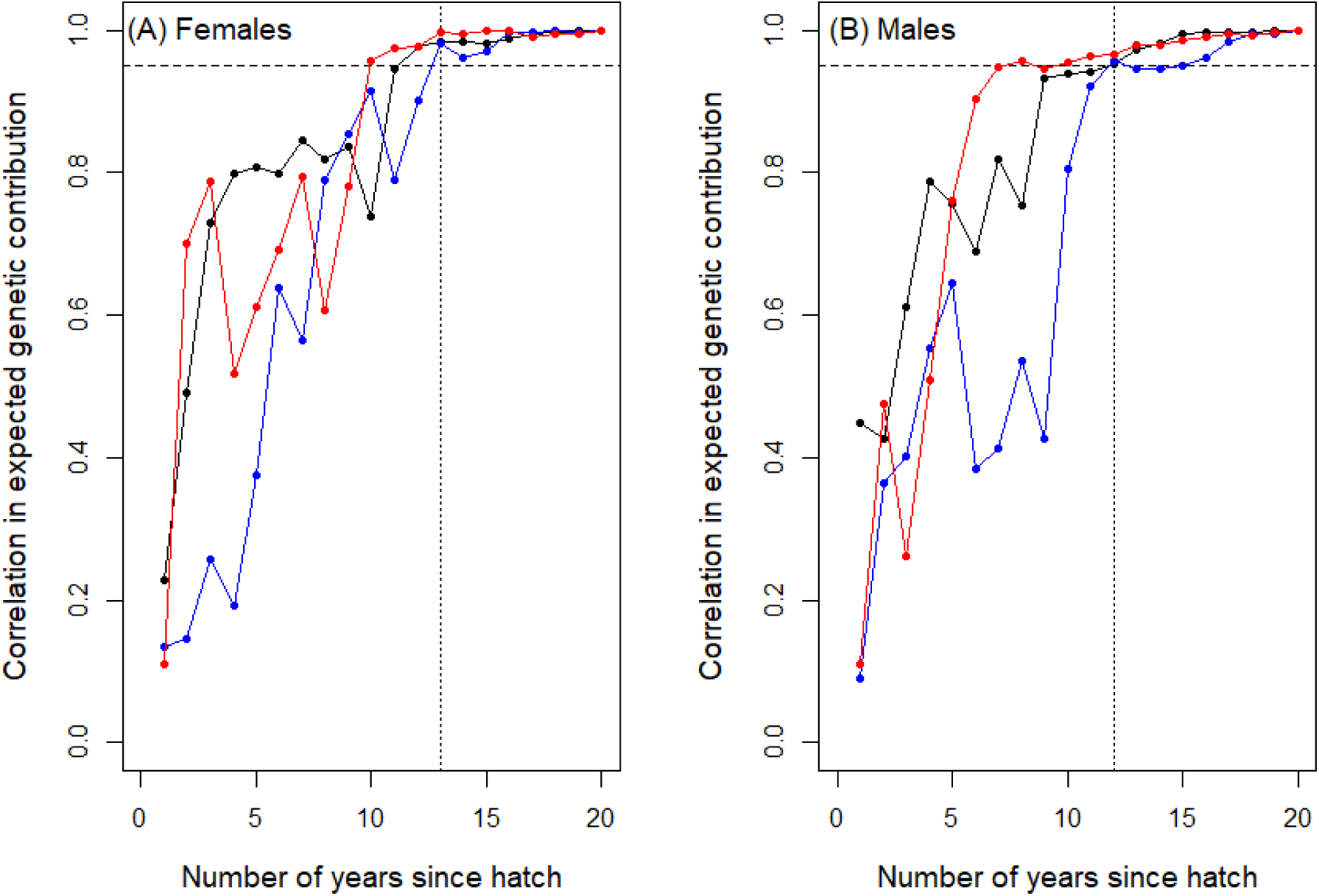
Pearson correlation coefficients between individuals’ absolute expected genetic contributions to the total extant population 20 years post-hatch versus each previous year for adult (A) female and (B) male songs sparrows hatched in 1992 (black), 1993 (blue) and 1994 (red) that survived to adulthood. Dashed horizontal lines demarcate correlations that equal or exceed 0.95. Dotted vertical lines demarcate the numbers of years post-hatch at which the correlation reached 0.95 for all three cohorts. Correlation coefficients were virtually identical given proportional rather than absolute expected genetic contributions.

The distributions of individuals’ expected genetic contributions (Figs. 1 & 2, Supporting Information S4) show that most females and males that survived to adulthood made zero contribution to the total extant population 20 years later (i.e. *V*_i_=0, Fig. 4). This is despite that many individuals had non-zero values for short-term metrics of fitness (e.g. adult LRS measured as ringed offspring, Figs. 1, 2 & 4). The sex-specific distributions of individual *V*_i_ were consequently highly skewed; few individuals per cohort contributed to the genetic composition of subsequent generations (Fig. 4). The variance in *V*_i_ was smaller than the variance in 0.5LRS in both sexes, both across individuals that survived to adulthood and across all hatched individuals, but the CV was slightly larger for *V*_i_ (Fig. 4).

**Figure 4.**
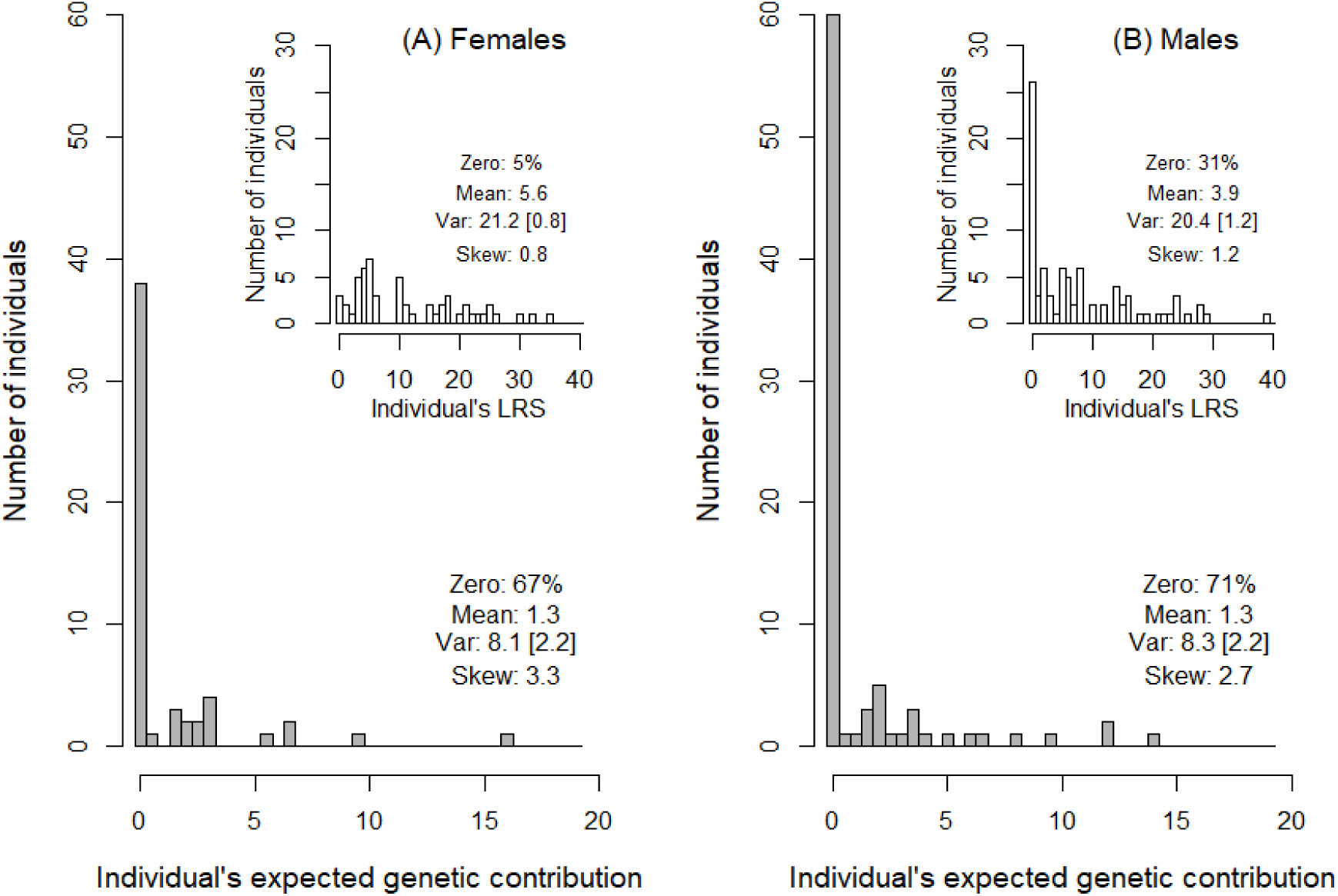
Distributions of individuals’ absolute expected genetic contributions to the total extant population 20 years post-hatch (i.e. estimated reproductive value, *V*_i_) in (A) 55 female and (B) 84 male song sparrows hatched during 1992-1994 that survived to adulthood and these individuals’ lifetime reproductive success (LRS) measured as ringed offspring (inset panels). Y-axis scales are standardised to facilitate comparison. Descriptive statistics comprise the percentage of individuals with values of zero, mean, variance (Var), skew and coefficient of variance (CV, in square brackets). Statistics were calculated for 0.5LRS to facilitate direct comparison with *V*_i_. Corresponding statistics for full distributions including values of zero for individuals that died before adulthood are: *V*_i_: mean 0.2 and 0.4, variance 1.7 and 2.8, skew 8.4 and 5.5, CV 5.6 and 4.7; and 0.5LRS: mean 1.0 and 1.2, variance 8.5 and 9.3, skew 3.3 and 3.1, CV 2.9 and 2.9, for females and males respectively. Further summary statistics are in Supporting Information S7.

Individual *V*_i_ estimated 20 years post-hatch tightly predicted the probability of allele extinction (*P*_E_) during the same timeframe (Fig. 5), as expected given the typically high *P*_E_ (Barton & Etheridge 2011). There was substantial variation in the number of allele copies present in the total extant population conditional on allele persistence (i.e. ≥1 copy), both among individuals (Fig. 6) and among gene-drop iterations within individuals (Supporting Information S6). The mean and variance in copy number were both strongly positively associated with individual *V*_i_, but the CV was independent of *V*_i_ on average (Fig. 6). This concurs with the expectation that, conditional on allele persistence, the distribution of genetic contributions will be independent of *V*_i_ (Barton & Etheridge 2011). The tight relationship between *V*_i_ and *P*_E_ that is already evident (Fig. 5) implies that *V*_i_ encapsulates an individual’s longer-term genetic contribution (Barton & Etheridge 2011).

**Figure 5.**
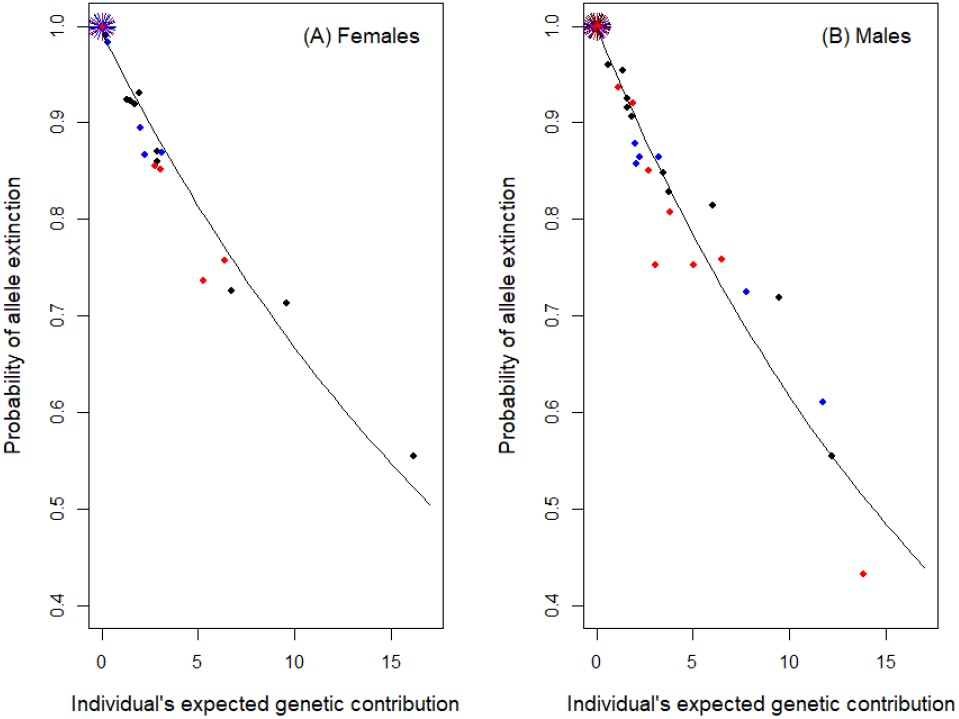
Relationships between an individual’s probability of allele extinction (*P*_E_) and absolute expected genetic contribution to the total extant population 20 years post-hatch (i.e. estimated reproductive value, *V*_i_) for (A) female and (B) male song sparrows hatched in 1992 (black), 1993 (blue) and 1994 (red) that survived to adulthood. Petals denote multiple individuals with zero expected genetic contribution, and hence *P*_E_=1. Solid lines depict the expected exponential relationship (i.e. *P*_E_ = exp(-α*V*_i_), Barton & Etheridge 2011) fitted to all three cohorts combined. Estimated regression coefficients α were −0.040 and −0.048 for females and males respectively for absolute *V*_i_, and −19.9 and −21.8 respectively for proportional *V*_i_.

**Figure 6.**
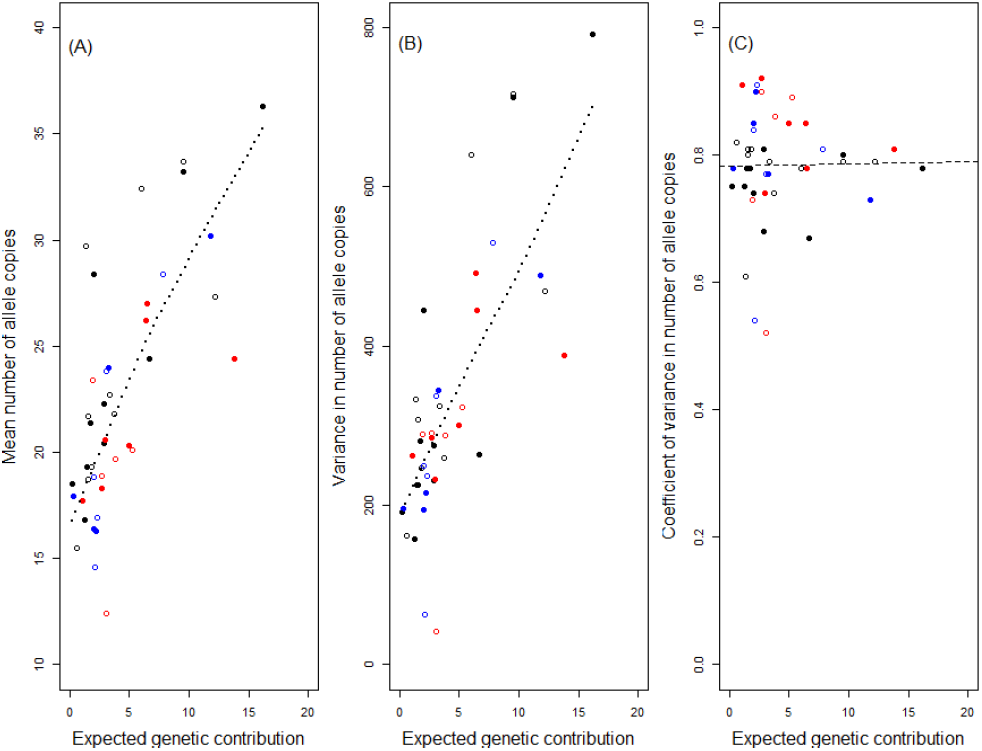
Relationships between an individual’s absolute expected genetic contribution to the total extant population 20 years post-hatch (i.e. estimated reproductive value, *V*_i_) and the (A) mean, (B) variance and (C) coefficient of variance in the number of allele copies conditional on allele persistence (i.e. ≥1 copy) for 18 female (filled symbols) and 24 male (open symbols) song sparrows hatched in 1992 (black), 1993 (blue) and 1994 (red) with *V*_i_ > 0. On A & B, dotted lines denote smoothed curves (regressions were not fitted because the theoretical form of relationship is unclear). On C, the dashed line denotes the linear regression (slope estimate: 0.001).

### Individual fitness metrics and reproductive value

Across the totals of 55 females and 84 males hatched during 1992-1994 that survived to adulthood, individual *V*_i_ estimated 20 years post-hatch was positively associated with each of the six metrics of short-term individual fitness (Figs. 7 & 8, Supporting Information S7). However, correlation coefficients were moderate for lifespan (∼0.3-0.4) and for metrics of ringed and independent offspring (∼0.4-0.5), and still far from unity for metrics of recruited offspring (∼0.6-0.7, Figs. 7 & 8, Supporting Information S7). There is considerable scatter around estimated linear regressions, meaning that short-term fitness metrics typically explain less than half the estimated among-individual variation in *V*_i_ (Figs. 7 & 8, Supporting Information S7). While the slopes of regressions of absolute *V*_i_ on absolute fitness metrics commonly diverged substantially from one, those for relative (i.e. mean-standardised) *V*_i_ on relative fitness metrics were fairly close to one (Figs. 7 & 8, Supporting Information S7). This implies that short-term metrics of individuals’ relative fitness can potentially provide unbiased predictors of relative genetic contributions to future generations on average, despite the considerable individual-level deviation. There was no substantial difference in predictive ability between LRS and λ_ind_ measured to analogous offspring life stages (Figs. 7 & 8, Supporting Information S7). These conclusions, and specifically regression slopes and R2 values, did not change markedly when including values of zero *V*_i_ and fitness for totals of 248 females and 194 males hatched during 1992-1994 that died before adulthood (Figs. 7 & 8, Supporting Information S7), or using proportional rather than absolute *V*_i_.

**Figure 7.**
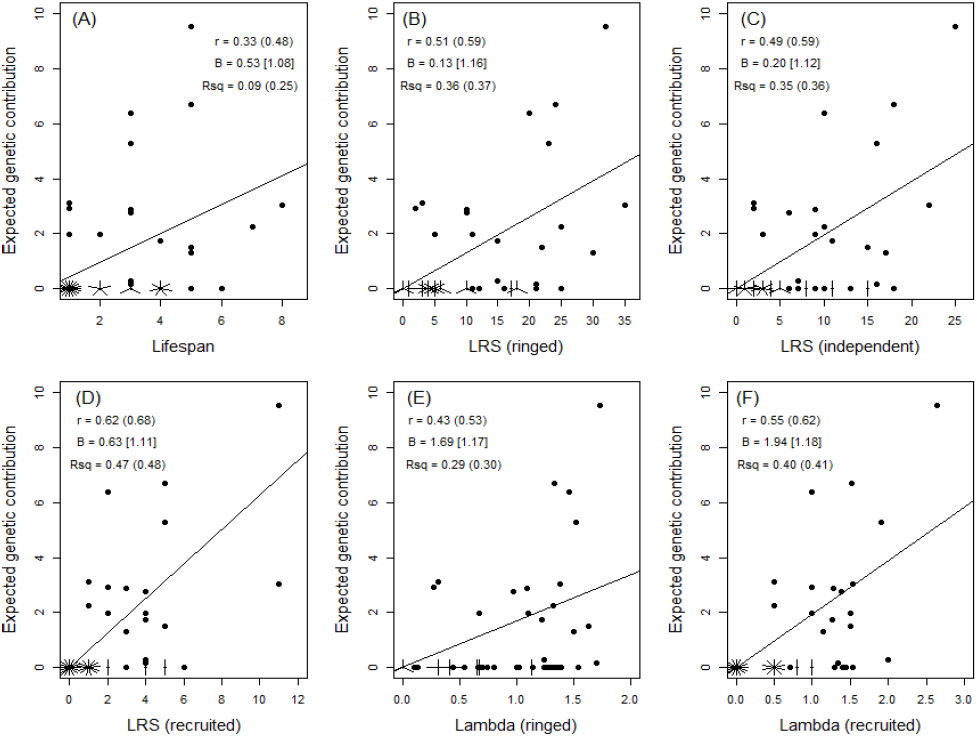
Relationships between an individual’s absolute expected genetic contribution to the total extant population 20 years post-hatch (i.e. estimated reproductive value, *V*_i_) and six short-term metrics of fitness across 55 female song sparrows hatched in 1992-1994 that survived to adulthood. Fitness metrics are (A) lifespan, lifetime reproductive success (LRS) measured as (B) ringed, (C) independent and (D) recruited offspring, and λ_ind_ (Lambda) measured across (E) ringed and (F) recruited offspring. Petals denote multiple individuals with identical values. Solid lines denote linear regressions (forced through the origin in B-F). Statistics are Pearson correlation coefficient (r), linear regression slope (B) and adjusted R2 (Rsq) calculated for absolute values of *V*_i_ and fitness across adults, or including individuals that died before adulthood (in parentheses), or using mean-standardised relative values (in square brackets). Further summary statistics are in Supporting Information S7.

**Figure 8.**
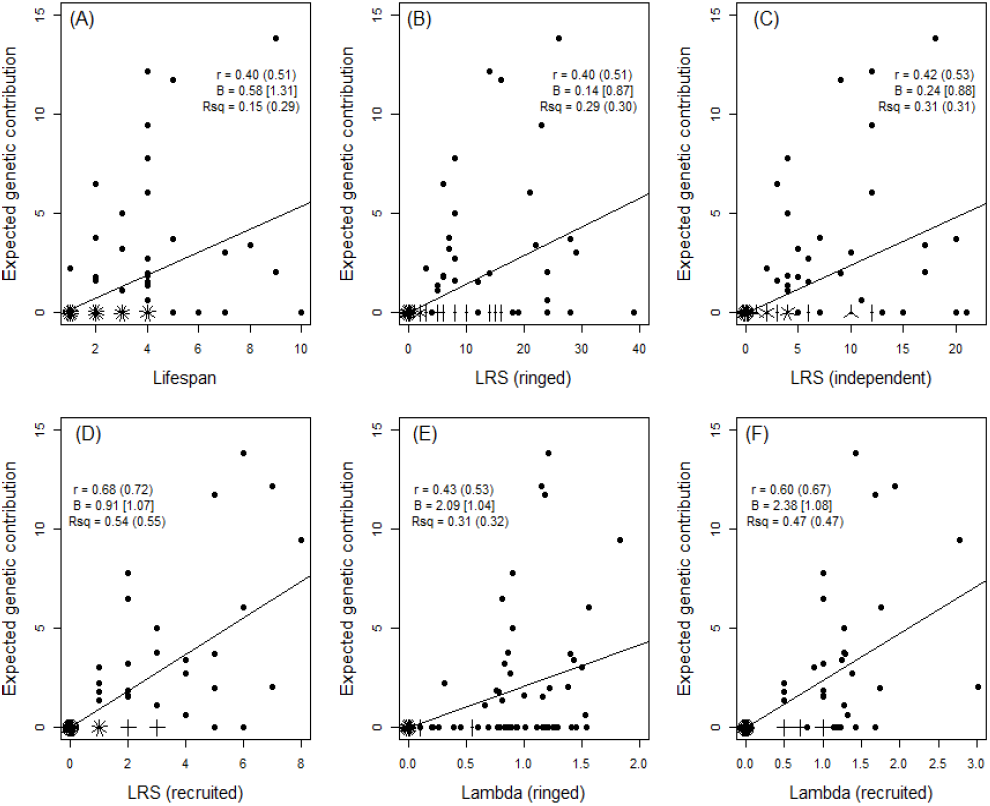
Relationships between an individual’s absolute expected genetic contribution to the total extant population 20 years post-hatch (i.e. estimated reproductive value, *V*_i_) and six short-term metrics of fitness across 84 male song sparrows hatched in 1992-1994 that survived to adulthood. Fitness metrics are (A) lifespan, lifetime reproductive success (LRS) measured as (B) ringed, (C) independent and (D) recruited offspring, and λ_ind_ measured across (E) ringed and (F) recruited offspring. Figure attributes are as for Figure 7. Further summary statistics are in Supporting Information S7.

## Discussion

All attempts to explain and predict evolutionary dynamics require some assessment of the fitness of alleles and/or individual organisms; yet theoretical fitness concepts can be hard to reconcile with empirical realities encompassing overlapping and varying generation times, environmental and demographic stochasticity, and context-dependent selection and genetic variation (de Jong 1994; Hunt & Hodgson 2010; Sæther & Engen 2015; Graves & Weinreich 2017). Relationships between short-term metrics of individual fitness and phenotypic trait values define selection gradients, which can be combined with information on additive genetic (co)variances to infer evolutionary outcomes (conditional on fulfilling key assumptions of evolutionary quantitative genetic theory, Lande & Arnold 1983). Yet, especially in systems where genetic (co)variances cannot be readily quantified and/or major cross-generation effects are postulated, evolutionary outcomes are widely discussed solely based on observed variation in individual or lineage fitness and associations with focal phenotypes (Kokko *et al.* 2003; Hunt *et al.* 2004; Hunt & Hodgson 2010). Any success of such approaches depends, not least, on the degree to which available short-term fitness metrics for individuals do reliably predict longer-term genetic contributions (Benton & Grant 2000; Brommer *et al.* 2002; Hunt *et al.* 2004). However, few studies have attempted to quantify such relationships in populations experiencing natural environmental, demographic, genetic and selective variation. Brommer *et al.* (2004) used partial pedigree data from open populations of collared flycatchers (*Ficedula albicollis*) and ural owls (*Strix uralensis*) to relate individual LRS and λ_ind_ to longer-term genetic contributions estimated over >2 generations. However, genetic contributions were only tracked through maternal lineages, meaning that persistence of autosomal alleles, which are propagated through both sexes, was probably greatly underestimated (Brommer *et al.* 2004). Individual LRS (measured as recruited offspring) and λ_ind_ explained only a third of observed variation in number of locally-fledged grand-offspring (i.e. one further generation) in long-tailed tits (*Aegithalos caudatus*, MacColl & Hatchwell 2004). Indeed, the general failure to measure multi-generation fitness has been highlighted as a short-coming in the context of testing sexual selection theory (Kokko *et al.* 2003; Hunt *et al.* 2004, but see Hunt & Hodgson 2010).

Our analyses of ≥20 years (i.e. ∼8 generations on average, Supporting Information S1 & S2) of locally complete, accurate, pedigree data from free-living song sparrows showed that individuals’ expected genetic contributions to the total extant population stabilised within the observed timeframe (Figs. 1-3). Individual reproductive value (*V*_i_, sensu Barton & Etheridge 2011) can consequently be estimated, and showed considerable among-individual variation in both sexes (Fig. 4). As expected, *V*_i_ tightly predicted the short-term probability of allele extinction (or, conversely, persistence), and is therefore likely to be directly informative regarding longer-term individual genetic contributions (Barton & Etheridge 2011). But, widely-used short-term metrics of individual fitness typically explained less than half the observed among-individual variation in *V*_i_ (Figs. 7 & 8).

The availability of tractable short-term metrics that do partially predict individuals’ expected longer-term genetic contributions in nature, despite the complexity and stochasticity of all underlying processes, could be deemed a success. However, there is unsurprisingly substantial unexplained variation and hence potential to draw erroneous evolutionary inferences directly from observed among-individual variation in short-term fitness. Discrepancies arise because some individuals with low (but non-zero) LRS and λ_ind_ made substantial longer-term genealogical and hence expected genetic contributions to the focal population, while many individuals with non-zero LRS and λ_ind_ made zero longer-term contributions (Figs. 1, 2 & 4; Supporting Information S4). These outcomes are not simply due to the small size of the focal population. Individual *V*_i_ results from Mendelian segregation of alleles conditional on an individual’s pedigree of descendants which, in sexually reproducing species, comprises biparental lineages (Chang 1999; Barton & Etheridge 2011). In species with limited reproductive capacity (e.g. any species with substantial parental care), probabilities of lineage and allele extinction will be high irrespective of total population size, simply due to individual-level environmental and demographic stochasticity and drift (e.g. Gravel & Steel 2015). However, lineage extinction probabilities must also depend on mean demographic rates and population dynamics (e.g. Metcalf & Parvard 2007; Gravel & Steel 2015; Graves & Weinreich 2017). The focal song sparrow population decreased in size in 1998-1999 (Supporting Information S1), which was sufficiently soon after the focal cohorts hatched to potentially eliminate all descendants of some individuals. Such lineage extinctions become less likely across subsequent generations, since all individuals with non-zero expected genetic contributions will, in the medium-term, be genealogical ancestors of all extant population members (Chang 1999; Caballero & Toro 2000; Barton & Etheridge 2011; but see Gravel & Steel 2015). However such variation in local population size, caused by environmentally-induced variation in fitness, will apply to almost all populations and sub-populations in nature, and will consequently be integral to any evolutionary outcome (Sæther & Engen 2015; Engen & Sæther 2017).

An individual’s observed LRS and λ_ind_ constitute stochastic realisations of its underlying propensity for fitness, which is not directly observable (McGraw & Caswell 1996; Link *et al.* 2002; Snyder & Ellner 2018). Yet, realised short-term fitness, comprising the number of offspring that was actually produced, might be envisaged as a reasonable predictor of longer-term genetic contribution that captures the implications of initial individual-level realisations of environmental and demographic stochasticity in survival and breeding success (e.g. Sæther & Engen 2015). Predictive capability will also depend partly on the additive genetic variance and heritability in LRS, and the latter is non-zero but small in song sparrows (<0.1 measured approximately chick-to-chick, Wolak *et al.* 2018). Such small or moderate values may be broadly typical, although still surprisingly few rigorous estimates are available (Shaw & Shaw 2014; Hendry *et al.* 2018). Consequently, the song sparrow data serve to illustrate the degree to which measures of longer-term evolutionary fitness are removed from short-term metrics of phenotypic fitness that are directly observable on individuals (e.g. de Jong 1994).

If direct estimation of individual *V*_i_ is, or soon could be, within reach of at least some field studies, what are its uses? *V*_i_ substantially provides the correct ‘answer’ as to which individuals are expected to make longer-term genetic contributions to any focal population, and should therefore represent a focus of adaptive evolution (Grafen 2006; Barton & Etheridge 2011). Yet, *V*_i_ does not fully describe the mean number of allele copies conditional on persistence (Fig. 6), which reflects the detailed structures of individuals’ pedigrees (Barton & Etheridge 2011; Gravel & Steel 2015). Further, *V*_i_ is phenomenological, and does not itself provide direct mechanistic insight into the processes or traits that cause long-term outcomes. Observed phenotypic variation of interest could potentially be related to *V*_i_ rather than to short-term fitness metrics, thereby better capturing associations between phenotypes and longer-term genetic contributions. However, since studies that can quantify individual *V*_i_ will necessarily have multi-generation pedigree (or genomic) data, such data might often be better deployed to estimate genetic covariances among traits of interest and metrics of fitness that span single generations or other short time intervals. In principle, and conditional on key quantitative genetic assumptions, such analyses can distinguish genetic from environmental covariances, quantify evolutionary constraints and cross-generational effects, and predict overall evolutionary outcomes (de Jong 1994; Morrissey *et al.* 2010; Reid 2012; Shaw & Shaw 2014). Yet, such approaches and multi-generation predictions also face considerable challenges, especially given density-, frequency- and environment-dependent selection and genetic variation, and inbreeding and other interactions among diverse relatives. Further consideration of the structure of individual pedigrees and genealogies and hence the basis of *V*_i_ might then provide useful complementary insights into age-specific contributions, lineage introgression, individuals responsible for inbreeding, and ultimately inclusive fitness (e.g. Caballero & Toro 2000; Suwanlee *et al.* 2007; Barton & Etheridge 2011; Newman & Easteal 2015).

## Supporting information

Reid_Supporting_Information

## Acknowledgements

We thank the Tsawout & Tseycum First Nations bands for access to Mandarte, everyone who contributed to long-term data collection, Ryan Germain for helpful comments, and the European Research Council, NSERC (Canada) and the Swiss National Science Foundation for funding. The authors declare no conflicts of interest.

## Author contributions

JMR conceived and undertook the analyses and drafted the manuscript. MEW wrote the gene-dropping algorithm. PA, LFK and PN undertook and oversaw field data collection and pedigree construction. All authors contributed to conceptual clarification and manuscript editing.

## Supporting information

**S1.** Study system, pedigree, population size and sex ratio.

**S2.** Generation time and adult ages.

**S3.** Metrics of individual fitness.

**S4.** Absolute expected genetic contributions for song sparrows hatched in 1992 and 1994.

**S5.** Proportional expected genetic contributions for song sparrows hatched in 1992-1994.

**S6.** Distributions of allele frequencies.

**S7.** Additional summary statistics.

