## Supplementary material for "Individuals’ expected genetic contributions to future generations, reproductive value, and short-term metrics of fitness in free-living song sparrows (Melospiza melodia)": Reid_Supporting_Information

### Supporting Information S1. Study system, pedigree, population size and sex ratio.

Song sparrows typically form socially monogamous breeding pairs with biparental care. Each year since 1975, all breeding attempts on Mandarte (typically 2-3 per pair per year) have been closely monitored, and all chicks surviving to six days post-hatch have been individually colour-ringed (Smith *et al.* 2006). Mandarte (approximately 6 hectares) lies within a large song sparrow meta-population and receives occasional immigrants (1.1 arrivals/year on average, constituting ~3% of new recruited adults) which are mist-netted and colour-ringed. All adults are therefore individually recognisable. Each spring, all surviving adults, and the social parents of all ringed chicks, are identified (Smith *et al.* 2006; Reid *et al.* 2014). All individuals ringed during 1993-2015 were then genotyped at ~160 polymorphic microsatellite markers, allowing true genetic parents to be assigned to all chicks with extremely high individual-level statistical confidence (>>0.99, Nietlisbach *et al.* 2015, 2017). Such analyses revealed zero extra-pair maternity but approximately 28% extra-pair paternity (Sardell *et al.* 2010; Reid *et al.* 2016), and yield complete genetically-verified pedigree data (Reid *et al.* 2014; Nietlisbach *et al.* 2017).

Adult song sparrows are highly philopatric, with probably little or no emigration. Local juvenile recruitment rates on Mandarte are also typically high: 24% of all hatched offspring recruited locally on average across years. Since mean survival from hatching to age one year is unlikely to be substantially higher, juvenile emigration must be relatively infrequent. Some emigration doubtless occurs, resulting in local loss of allele copies. However, the observed pedigree allows calculation of expected relatedness and any desired metric of fitness across adults and ringed chicks with most likely zero error or missing data with respect to the local Mandarte population (Reid *et al.* 2014, 2016; Wolak *et al.* 2018). It therefore provides a complete depiction of local fitness and expected local allele persistence.

The defined total extant population in each year, and hence the annual allele totals, exclude allele copies inherited by zygotes that died before six days old (i.e. the age of ringing and genetic parentage assignment). However, this exclusion does not impede the current objective of evaluating adults' stabilised $V_i$ because dead zygotes cannot directly contribute to the population's genetic composition in subsequent years or generations.

Estimates of $V_i$ for individuals hatched in 1992-1994 are partially non-independent; some individual pedigrees are nested because some individuals hatched in 1993-1994 are the offspring of individuals hatched in 1992-1993. However, because of the substantially overlapping generations, some parents hatched before 1992 contributed offspring to two or three cohorts, meaning that these individuals' pedigrees of descendants are not nested with respect to the current analyses.

All song sparrow chicks hatched in 1993 and 1994 were sexed by genotyping the chromobox-helicase-DNA-binding (CHD) gene (Postma *et al.* 2011). Chicks hatched in 1992 were not genotyped, but sexes of individuals that survived to adulthood were ascertained from reproductive behaviour, and numbers of females and males that did not survive to adulthood were estimated assuming a primary cohort sex ratio of 1:1 across ringed chicks (as observed on average across subsequent years, Postma *et al.* 2011).

**Figure S1.** Numbers of song sparrows (*Melospiza melodia*) alive in the focal Mandarte island population in each year during 1975-2015, showing adult (≥1 year old) females (black circles), adult males (mid-grey circles), ringed chicks (light grey circles) and all individuals combined (black squares and dashed lines). Mid-grey shading highlights the natal years of the three focal cohorts (1992-1994). Light grey shading highlights the period (1993-2015) for which all parentage was genetically assigned and hence the population pedigree is known without error, allowing assessment of individuals' expected genetic contributions to the defined population. Note that the adult sex ratio is commonly male biased. Data are incomplete for 1981 due to reduced fieldwork in that year. Summary statistics for total adult population size through the focal period of 1992-2015: arithmetic mean = 73.3, range 33-128, geometric mean

= 68.2, harmonic mean = 63.4. The substantial among individual variance in lifetime reproductive success (e.g. Fig. 4) means that the local effective population size ($N_e$) will be smaller than census population size (O'Connor *et al.* 2006).

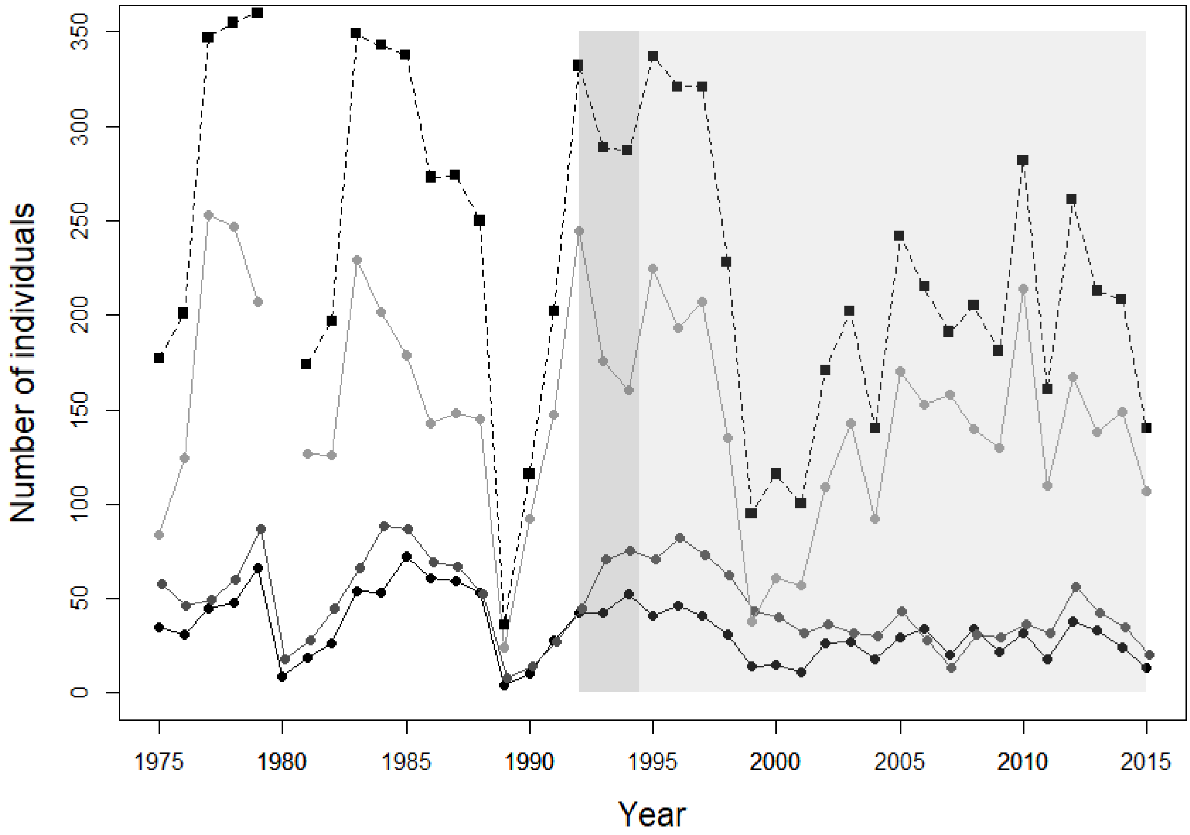

### Supporting Information S2. Generation time and adult ages.

Song sparrow generation time estimated as the mean age of parents across the focal study years (Fig. S2), and summary of the distribution of observed adult ages (Fig. S3).

**Figure S2.** Mean ages of mothers (black circles) and fathers (grey circles) of song sparrow chicks hatched on Mandarte in each year during 1992-2015. Mean parent age was 2.47±1.46SD across the grand total of 6954 observed parent ages (black line). Mean ages of mothers and fathers were 2.25±1.36SD (dashed line) and 2.70±1.52SD (grey line) respectively. Additionally, the mean ages of mothers and fathers of chicks that survived to adulthood (age one year) were 2.29±1.35SD and 2.77±1.63SD respectively (mean 2.53±1.51SD across the grand total of 1288 observations). These overall means capture the essence of the song sparrow life-history and hence provide a useful summary metric, but hide considerable among-year variation in mean parent age (as shown) and hence variation in generation time.

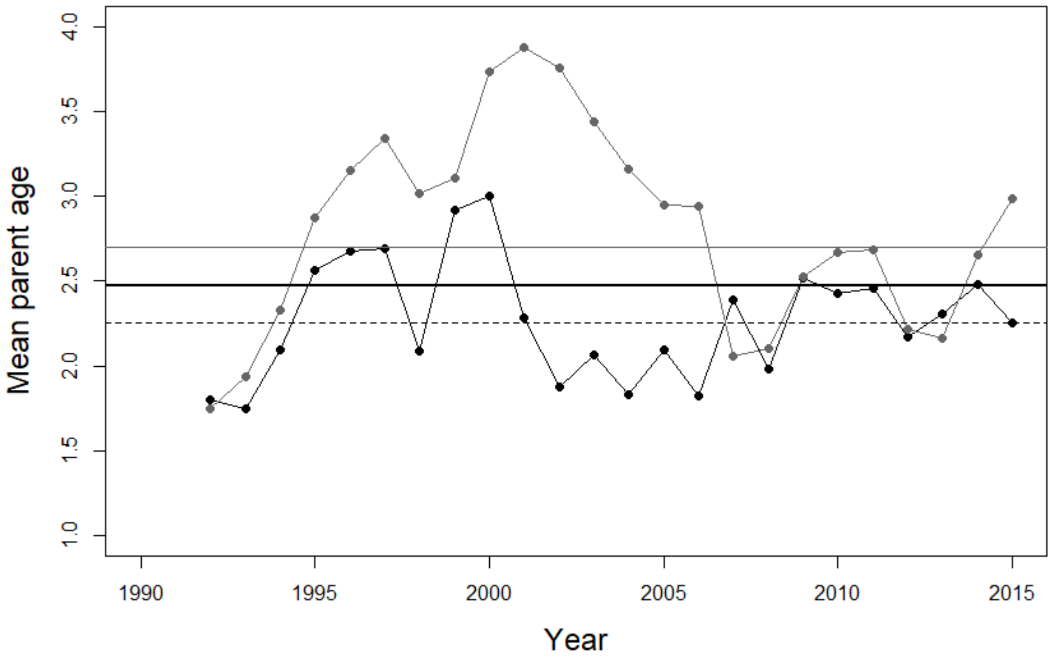


**Figure S3.** Frequency distribution of ages of adult song sparrows alive in 1992-2015. Mean adult age across the grand total of 1761 individual adult-years was 2.34±1.59SD. Mean ages of adult females and males were 2.18±1.44SD and 2.45±1.68SD respectively.

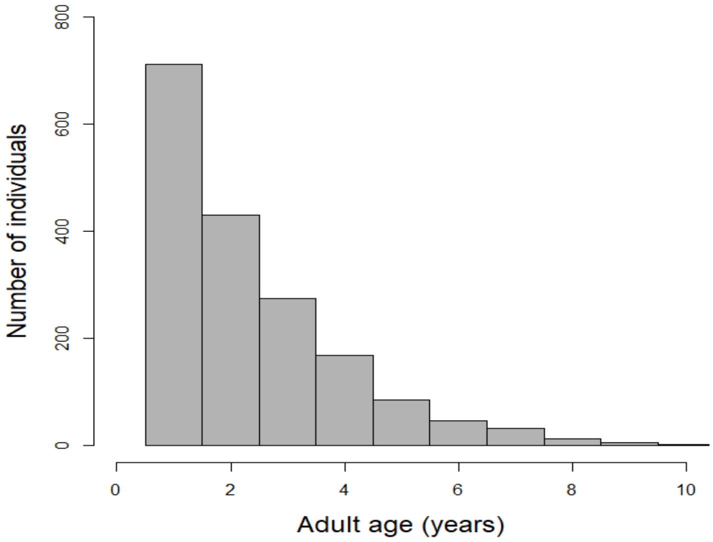

### Supporting Information S3. Metrics of individual fitness.

**Details of individual projection matrices and $\lambda_{ind}$.**

Individual projection matrices were formulated according to pre-breeding censuses for each individual song sparrow from the focal cohorts that survived to age one year (Table S1).

**Table S1.** Fully age-structured individual projection matrix formulated with a pre-breeding census. $F_1$ is an individual's fecundity aged one year, and $F_d$ is an individual's fecundity in its final breeding season before death. Subdiagonal values of 1 denote that an individual was observed to survive to the next year. The pre-breeding census formulation allows these elements to be enumerated at the level of the focal individual.

| | | | |
|---|---|---|---|
| $F_1$ | … | $F_{d-1}$ | $F_d$ |
| 1 | … | 0 | 0 |
| … | … | … | … |
| 0 | … | 1 | 0 |

Two values of $\lambda_{ind}$ were calculated for each individual, as the dominant eigenvalues of matrices with top row fecundity terms specified as:

1) $0.5.m_{ring}.\phi_j$, where $m_{ring}$ is the number of ringed offspring produced by each focal individual at each age and $\phi_j$ is the mean population-wide juvenile survival probability (i.e. ringing to age one year) in the focal year.

2) $0.5.m_{rec}$, where $m_{rec}$ is the number of recruited offspring produced by each focal individual at each age.

In both cases, the factor of 0.5 accounts for the transmission probability of a focal parental allele to each offspring given Mendelian inheritance.

As expected, measures of $\lambda_{ind}$ were strongly positively correlated with measures of lifetime reproductive success (LRS) across adults (Fig. S4). Values of both LRS and $\lambda_{ind}$ are zero for individuals that did not survive to adulthood (age one year).

**Figure S4.** Illustrative relationships between individual lifetime reproductive success (LRS) and $\lambda_{ind}$ for (A & B) 55 adult female and (C & D) 84 adult male song sparrows, with LRS and $\lambda_{ind}$ measured to (A & C) ringed chicks and (B & D) recruited offspring respectively. Petals denote multiple individuals with identical values.

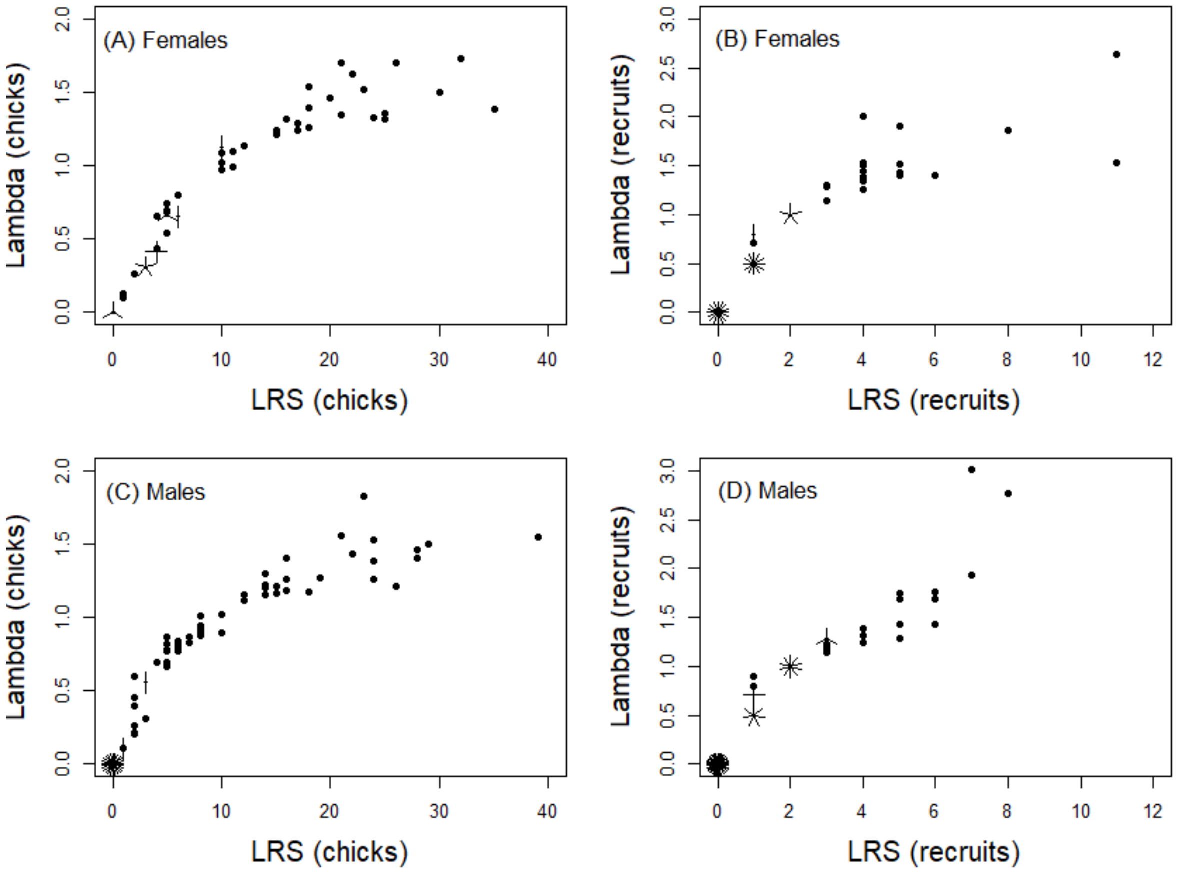

Numerous alternative metrics of individual fitness have been defined for use in wild populations with individual-based data on survival and reproduction, serving different purposes (e.g. Coulson *et al.* 2006; Moorad 2014; Sæther & Engen 2015). However these metrics are not considered here: Moorad's (2014) metric assumes no variation in demography or hence population growth rate within cohorts' lifespans (which is substantially violated in song sparrows); Coulson *et al.*'s (2006) and Sæther & Engen's (2015) metrics are primarily designed to consider annual fitness (measured adult-to-adult) rather than lifetime fitness.

### Supporting Information S4. Absolute expected genetic contributions for female and male song sparrows hatched in 1992 and 1994.

**Figure S5.** Observed lifetime reproductive success (LRS) measured as ringed offspring (top row) and absolute expected genetic contributions of 21 individual female song sparrows hatched in 1992 that survived to adulthood (columns) to the total extant population 1-20 years post-hatch (descending rows). Black shading denotes genetic contributions arising because a focal female was still alive in the focal year. Dark grey, light grey and white shading denote expected genetic contributions to offspring produced in the focal year, to surviving offspring produced in previous years, and to all subsequent descendants respectively. All bars (except LRS) are scaled to maximum y-axis values of 12 allele copies to allow direct comparison across years. Columns (i.e. females) are ordered by increasing expected contributions across final observed years.

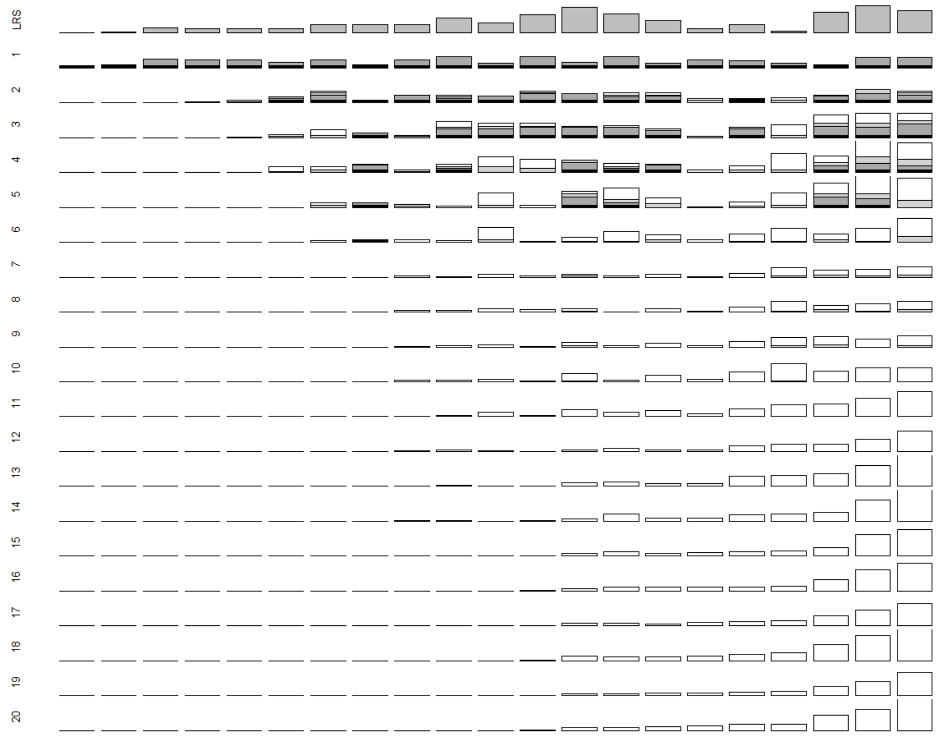

**Figure S6.** Observed lifetime reproductive success (LRS) measured as ringed offspring (top row) and absolute expected genetic contributions of 38 individual male song sparrows hatched in 1992 that survived to adulthood (columns) to the total extant population 1-20 years post-hatch (descending rows). Black shading denotes genetic contributions arising because a focal male was still alive in the focal year. Dark grey, light grey and white shading denote expected genetic contributions to offspring produced in the focal year, to surviving offspring produced in previous years, and to all subsequent descendants respectively. All bars (except LRS) are scaled to maximum y-axis values of 12 allele copies to allow direct comparison across years. Columns (i.e. males) are ordered by increasing expected contributions across final observed years.

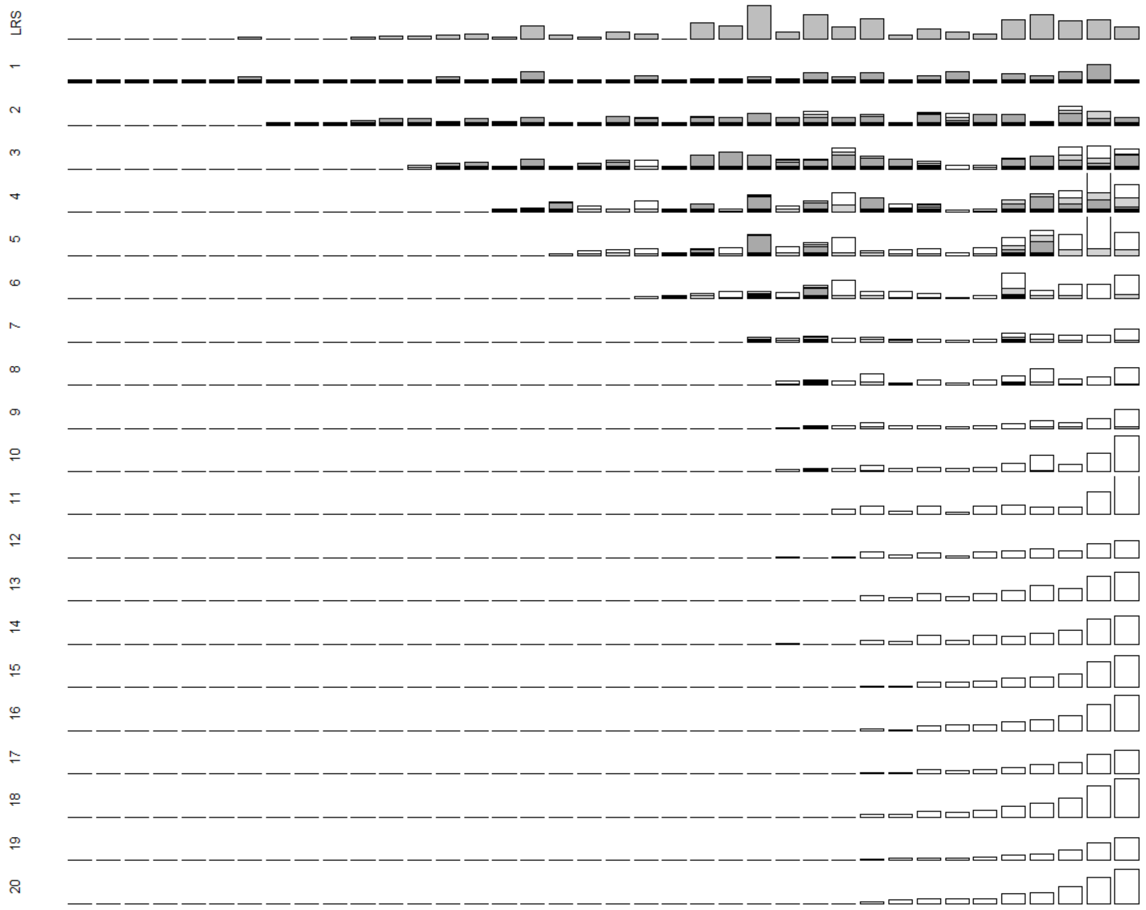

**Figure S7.** Observed lifetime reproductive success (LRS) measured as ringed offspring (top row) and absolute expected genetic contributions of 10 individual female song sparrows hatched in 1994 that survived to adulthood (columns) to the total extant population 1-20 years post-hatch (descending rows). Black shading denotes genetic contributions arising because a focal female was still alive in the focal year. Dark grey, light grey and white shading denote expected genetic contributions to offspring produced in the focal year, to surviving offspring produced in previous years, and to all subsequent descendants respectively. All bars (except LRS) are scaled to maximum y-axis values of 10 allele copies to allow direct comparison across years. Columns (i.e. females) are ordered by increasing expected contributions across final observed years.

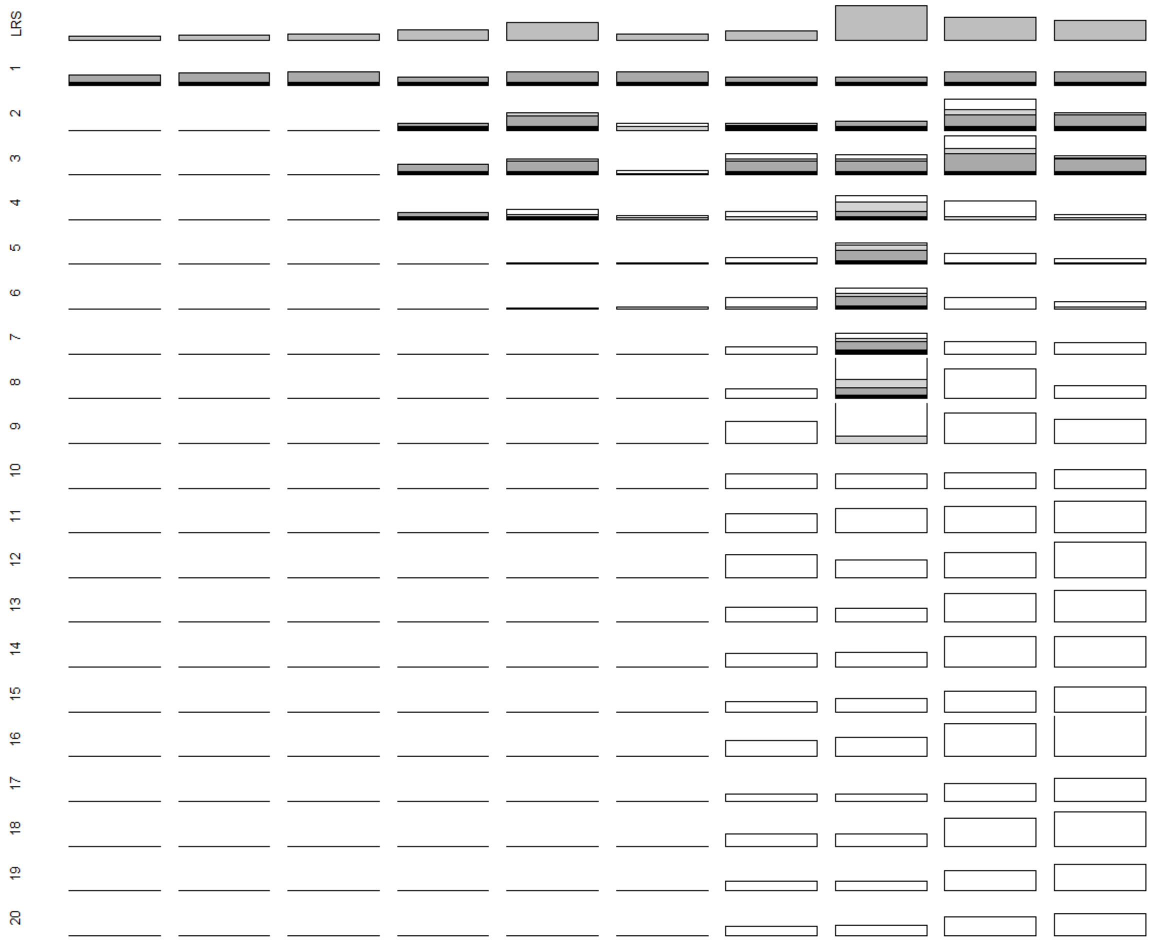

**Figure S8.** Observed lifetime reproductive success (LRS) measured as ringed offspring (top row) and absolute expected genetic contributions of 23 individual male song sparrows hatched in 1994 that survived to adulthood (columns) to the total extant population 1-20 years post-hatch (descending rows). Black shading denotes genetic contributions arising because a focal male was still alive in the focal year. Dark grey, light grey and white shading denote expected genetic contributions to offspring produced in the focal year, to surviving offspring produced in previous years, and to all subsequent descendants respectively. All bars (except LRS) are scaled to maximum y-axis values of 10 allele copies to allow direct comparison across years. Columns (i.e. males) are ordered by increasing expected contributions across final observed years. Values are truncated for one male with substantial expected genetic contributions (right column).

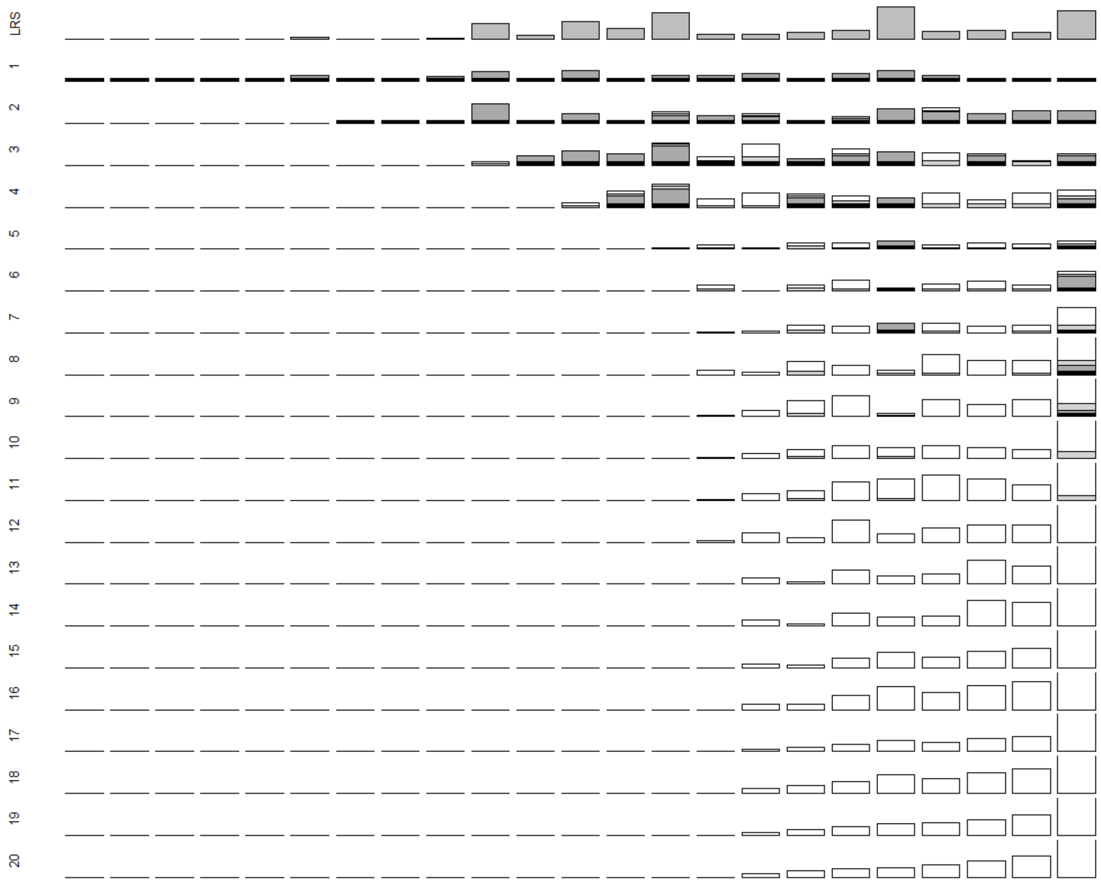

### Supporting Information S5. Proportional expected genetic contributions for female and male song sparrows hatched in 1992-1994.

**Figure S9.** Proportional expected genetic contributions of 21 individual female song sparrows hatched in 1992 that survived to adulthood (columns) to the total extant population 1-20 years post-hatch (descending rows). All bars are scaled to a maximum y-axis value of 0.03 to allow direct comparison across years. Columns (i.e. females) are ordered by increasing expected contributions across final observed years.

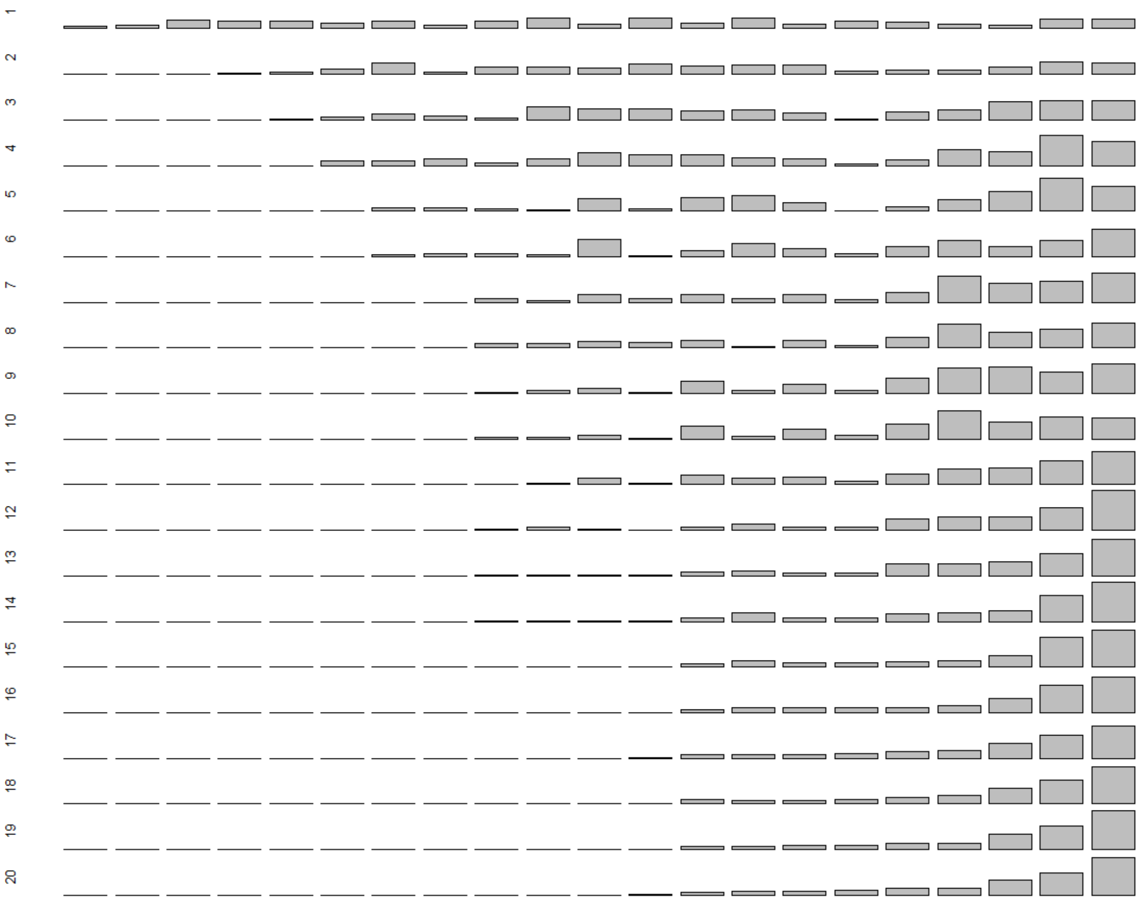

**Figure S10.** Proportional expected genetic contributions of 38 individual male song sparrows hatched in 1992 that survived to adulthood (columns) to the total extant population 1-20 years post-hatch (descending rows). All bars are scaled to a maximum y-axis value of 0.03 to allow direct comparison across years. Columns (i.e. males) are ordered by increasing expected contributions across final observed years.

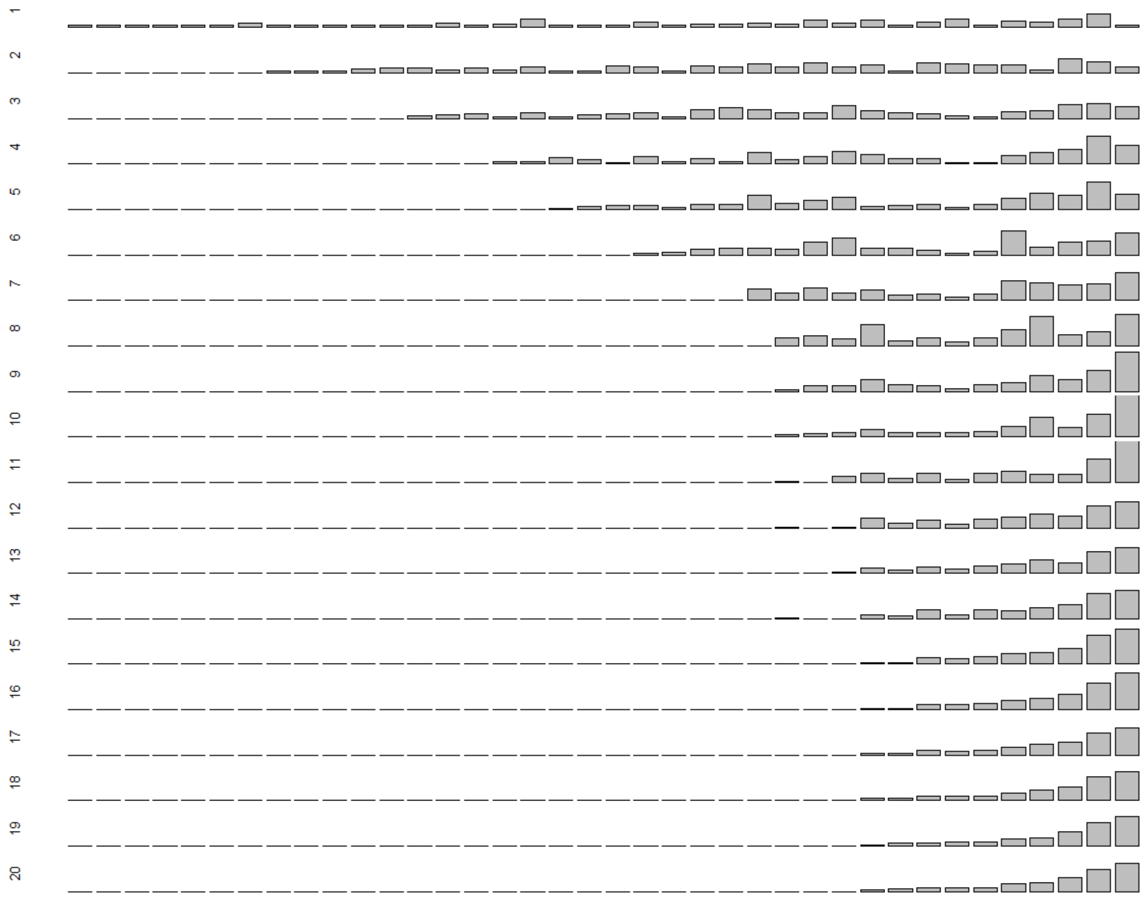

**Figure S11.** Proportional expected genetic contributions of 24 individual female song sparrows hatched in 1993 that survived to adulthood (columns) to the total extant population 1-20 years post-hatch (descending rows). All bars are scaled to a maximum y-axis value of 0.02 to allow direct comparison across years. Columns (i.e. females) are ordered by increasing expected contributions across final observed years.

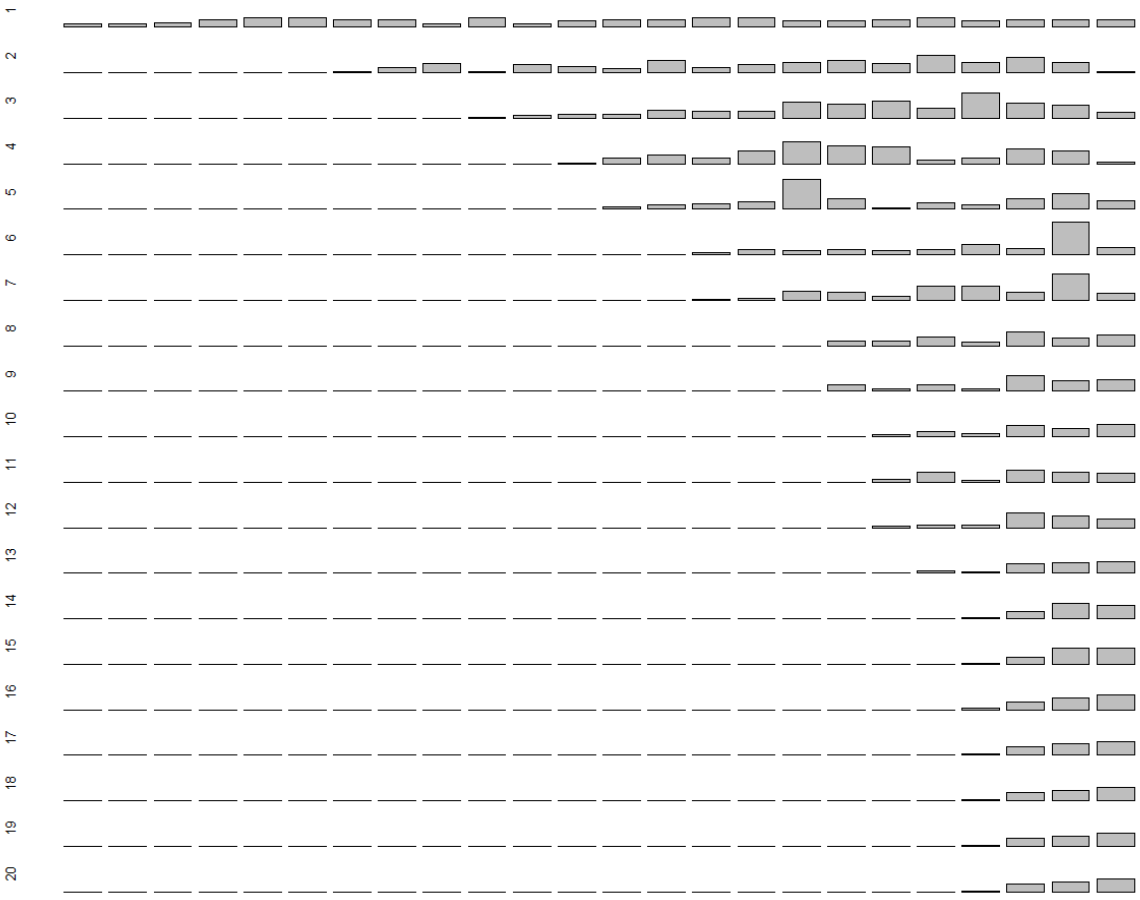

**Figure S12.** Proportional expected genetic contributions of 23 individual male song sparrows hatched in 1993 that survived to adulthood (columns) to the total extant population 1-20 years post-hatch (descending rows). All bars are scaled to a maximum y-axis value of 0.025 to allow direct comparison across years. Columns (i.e. males) are ordered by increasing expected contributions across final observed years.

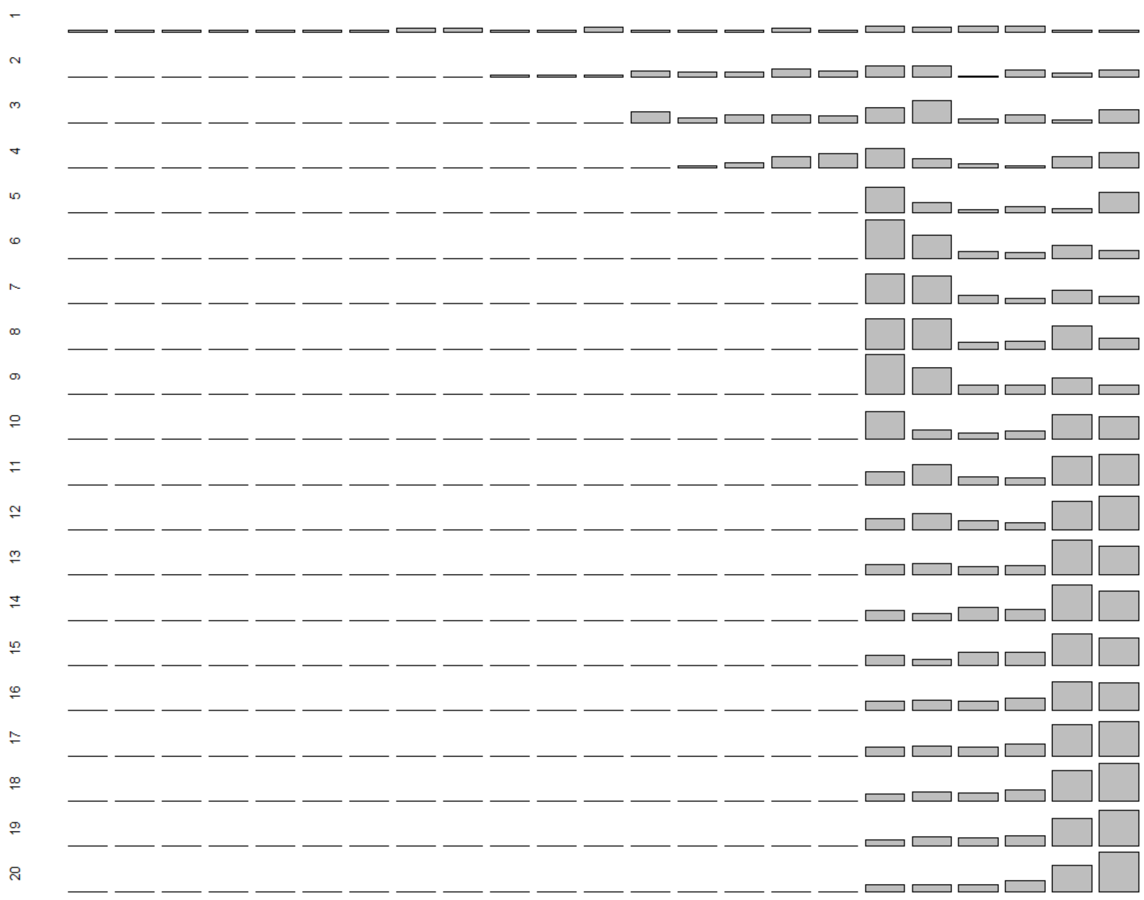

**Figure S13.** Proportional expected genetic contributions of 10 individual female song sparrows hatched in 1994 that survived to adulthood (columns) to the total extant population 1-20 years post-hatch (descending rows). All bars are scaled to a maximum y-axis value of 0.025 to allow direct comparison across years. Columns (i.e. females) are ordered by increasing expected contributions across final observed years.

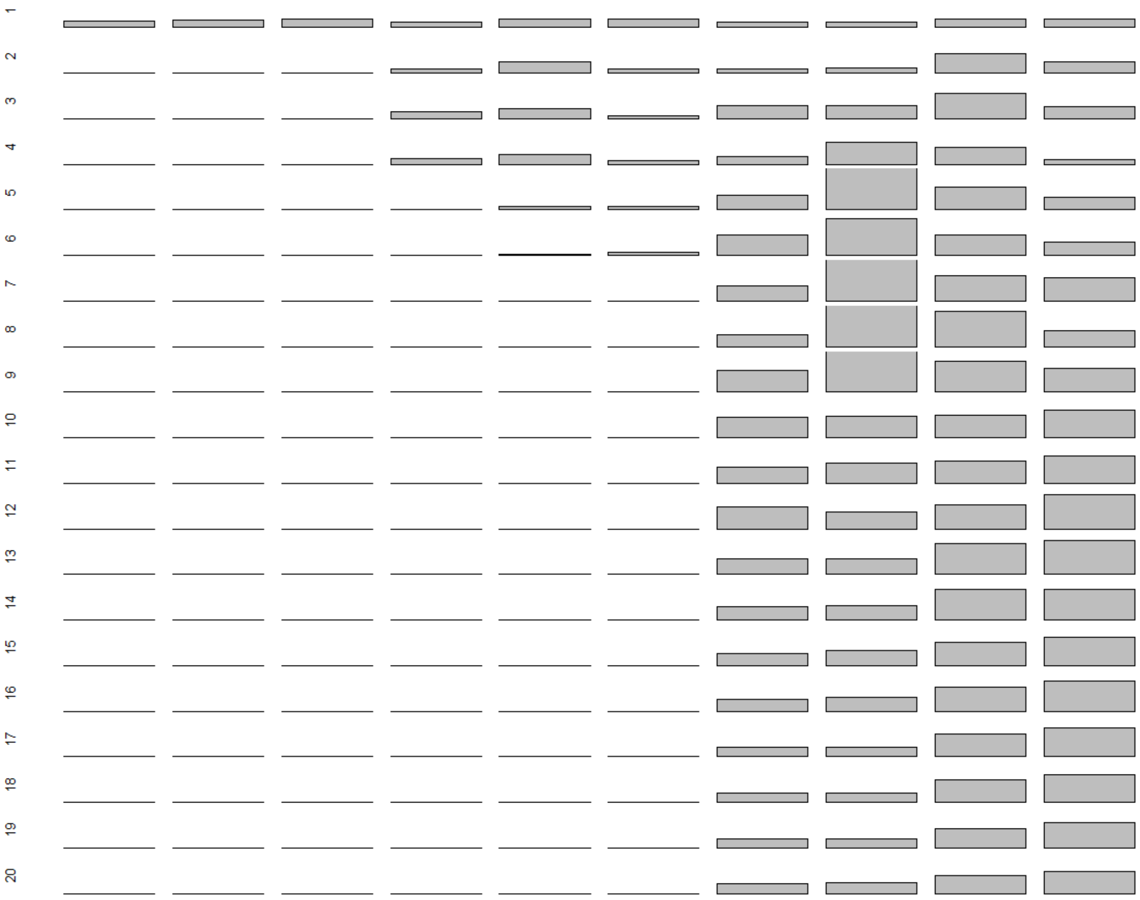

**Figure S14.** Proportional expected genetic contributions of 23 individual male song sparrows hatched in 1994 that survived to adulthood (columns) to the total extant population 1-20 years post-hatch (descending rows). All bars are scaled to a maximum y-axis value of 0.03 to allow direct comparison across years. Columns (i.e. males) are ordered by increasing expected contributions across final observed years. Values are truncated for one male with substantial expected genetic contributions (right column).

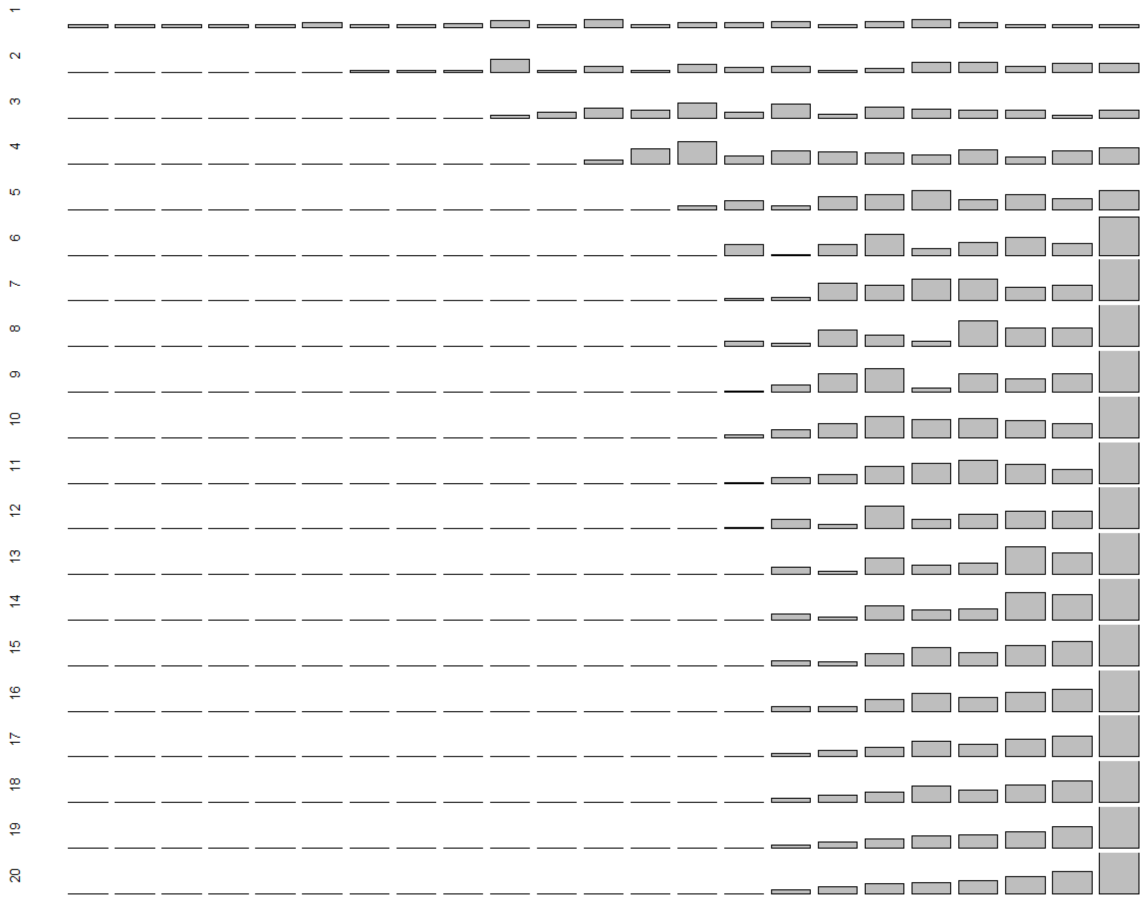

### Supporting Information S6. Distributions of allele frequencies.

**Figure S15.** Example frequency distributions of the number of allele copies present in the total extant population 20 years post-hatch for four adult female (dark grey bars) and six adult male (white bars) song sparrows hatched in 1993 with reproductive values (i.e. expected genetic contributions) greater than zero (see Figs. 1 & 2). Distributions were computed across 8,000 gene-drop iterations. For all individuals, the probability of allele extinction (P(E)) was high, and the associated high frequency of zero allele copies is not depicted. The mean, variance (Var), skew and coefficient of variance (CV=standard deviation/mean) in allele copy number conditional on persistence are shown for each individual. Y-axis scales are standardised to facilitate comparison among individuals.

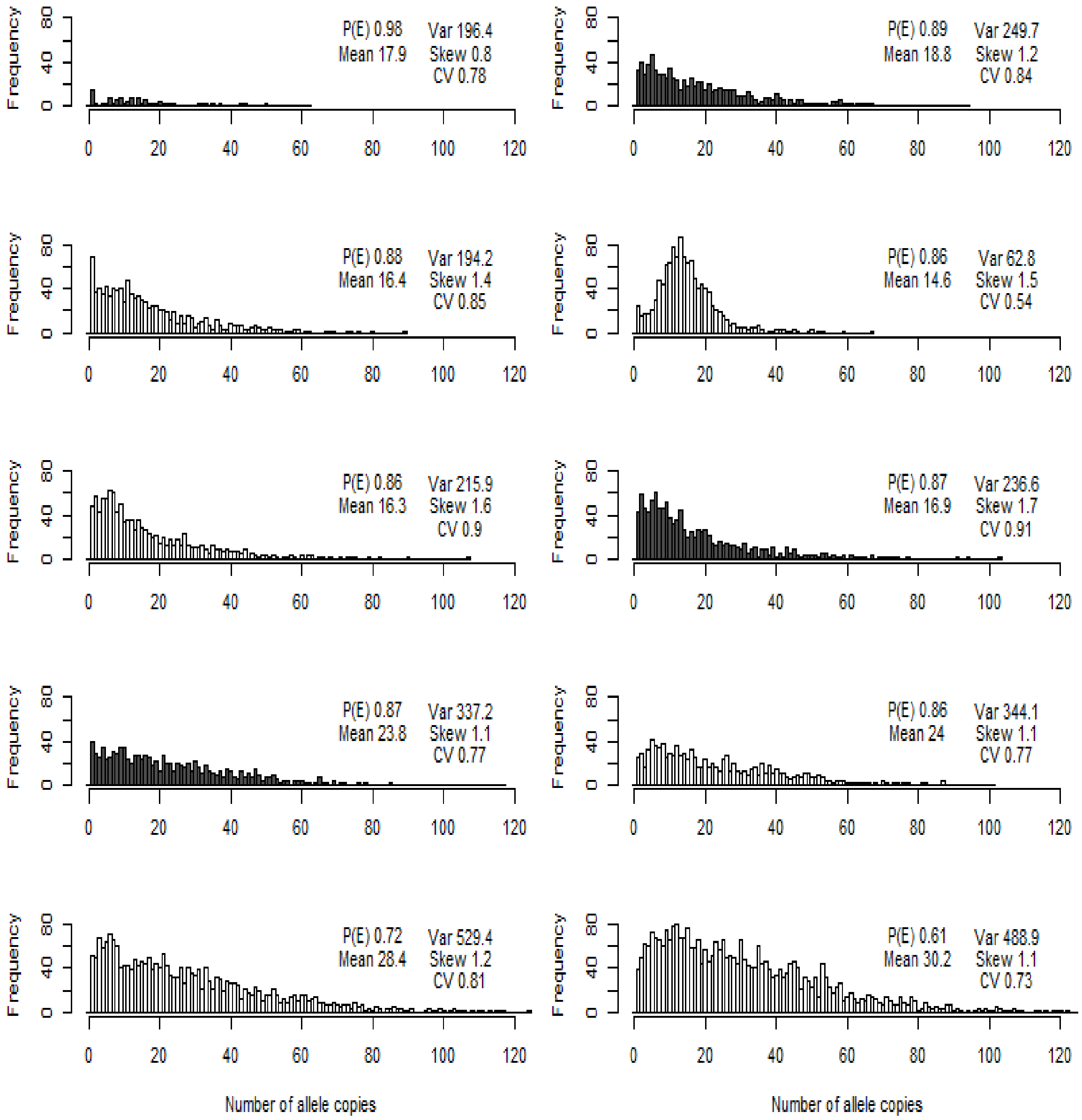
Frequency
P(E) 0.98
Mean 17.9
Var 196.4
Skew 0.8
CV 0.78
P(E) 0.89
Mean 18.8
Var 249.7
Skew 1.2
CV 0.84
P(E) 0.88
Mean 16.4
Var 194.2
Skew 1.4
CV 0.85
P(E) 0.86
Mean 14.6
Var 62.8
Skew 1.5
CV 0.54
P(E) 0.86
Mean 16.3
Var 215.9
Skew 1.6
CV 0.9
P(E) 0.87
Mean 16.9
Var 236.6
Skew 1.7
CV 0.91
P(E) 0.87
Mean 23.8
Var 337.2
Skew 1.1
CV 0.77
P(E) 0.86
Mean 24
Var 344.1
Skew 1.1
CV 0.77
P(E) 0.72
Mean 28.4
Var 529.4
Skew 1.2
CV 0.81
P(E) 0.61
Mean 30.2
Var 488.9
Skew 1.1
CV 0.73
Number of allele copies

### Supporting Information S7. Additional summary statistics.

**Table S1.** Summary statistics describing the distributions of six metrics of fitness, comprising individuals' lifespan, lifetime reproductive success (LRS) measured to ringed, independent (indep.) or recruited offspring, and $\lambda_{ind}$ measured to ringed or recruited offspring for (A) female and (B) male song sparrows. Statistics are calculated across 55 females and 84 males hatched in 1992-1994 that survived to adulthood (age one year) on Mandarte, and recalculated including values of zero for totals of 248 females and 194 males hatched in these cohorts that did not survive to adulthood (values in parentheses). Statistics are the mean, variance (Var), coefficient of variance (CV=standard deviation/mean) and skew. The coefficient of variance squared equals the opportunity for selection ($I$=variance/mean$^2$).

| | | Lifespan | LRS – ringed | LRS – indep. | LRS – recruited | $\lambda_{ind}$ – ringed | $\lambda_{ind}$ – recruited |
|---|---|---|---|---|---|---|---|
| (A) Females | Mean | 2.6 (0.5) | 5.6 (1.0) | 3.6 (0.7) | 1.1 (0.2) | 0.9 (0.2) | 0.8 (0.1) |
| | Var | 3.1 (1.6) | 21.2 (8.5) | 9.8 (3.7) | 1.7 (0.5) | 0.3 (0.2) | 0.4 (0.2) |
| | CV | 0.7 (2.6) | 0.8 (2.9) | 0.9 (2.9) | 1.2 (3.4) | 0.6 (2.5) | 0.9 (2.9) |
| | Skew | 0.9 (3.1) | 0.7 (3.3) | 0.8 (3.4) | 1.6 (4.7) | -0.1 (2.5) | 0.4 (3.1) |
| (B) Males | Mean | 3.0 (0.7) | 3.9 (1.0) | 2.4 (0.6) | 0.8 (0.2) | 0.6 (0.2) | 0.6 (0.1) |
| | Var | 3.9 (2.6) | 20.4 (8.0) | 7.9 (3.0) | 1.0 (0.4) | 0.3 (0.2) | 0.5 (0.2) |
| | CV | 0.7 (2.2) | 1.2 (2.9) | 1.2 (2.9) | 1.4 (3.2) | 0.9 (2.4) | 1.2 (2.9) |
| | Skew | 1.4 (2.7) | 1.2 (3.5) | 1.2 (3.5) | 1.4 (3.8) | 0.1 (2.3) | 1.1 (3.3) |

**Table S2.** Summary statistics describing relationships between individuals' expected genetic contributions to the total extant population 20 years post-hatch, interpreted as individual reproductive value ($V_i$), and six short-term metrics of individual fitness for (A) female and (B) male song sparrows hatched in 1992-1994. The six fitness metrics are individual lifespan, lifetime reproductive success (LRS) measured to ringed, independent (indep.) and recruited offspring, and $\lambda_{ind}$ measured to ringed and recruited offspring. Summary statistics are (i) Pearson correlation coefficient ($r_p$), (ii) Spearman correlation coefficient ($r_s$), (iii) linear regression slope (B), and (iv) adjusted $R^2$ (adj $R^2$). These statistics were calculated across 55 females and 84 males that survived to adulthood, with regressions forced through the origin (except for lifespan), since individuals with zero LRS and $\lambda_{ind}$ must have zero $V_i$. Statistics were then recalculated including 248 females and 194 males that died before adulthood, with all regressions forced through the origin (values in parentheses). Values of B were identical with and without the extra individuals, except for lifespan. Regression slopes were additionally calculated using relative $V_i$ and fitness metrics (i.e. individual value divided by the mean across all same-sex individuals, values in square brackets), and were again identical with and without the extra individuals, except for lifespan. Correlation coefficients and adjusted $R^2$ values are the same for 0.5LRS as for LRS, but regression slopes should be multiplied by two. $R^2$ values differ from $r_p^2$ because regressions were forced through the origin.

| | | Lifespan | LRS – ringed | LRS – indep. | LRS – recruited | $\lambda_{ind}$ – ringed | $\lambda_{ind}$ – recruited |
|---|---|---|---|---|---|---|---|
| A) Females | i) $r_p$ | 0.33<br>(0.48) | 0.51<br>(0.59) | 0.49<br>(0.59) | 0.62<br>(0.68) | 0.43<br>(0.53) | 0.55<br>(0.62) |
| | ii) $r_s$ | 0.40<br>(0.57) | 0.48<br>(0.59) | 0.48<br>(0.60) | 0.62<br>(0.68) | 0.46<br>(0.59) | 0.61<br>(0.68) |
| | iii) B | 0.53<br>[1.08]<br>(0.50)<br>[1.03] | 0.13<br>[1.16]<br>(0.13)<br>[1.16] | 0.20<br>[1.12]<br>(0.20)<br>[1.12] | 0.63<br>[1.11]<br>(0.63)<br>[1.11] | 1.69<br>[1.17]<br>(1.69)<br>[1.17] | 1.94<br>[1.18]<br>(1.94)<br>[1.18] |
| | iv) adj $R^2$ | 0.09<br>(0.25) | 0.36<br>(0.37) | 0.35<br>(0.36) | 0.47<br>(0.48) | 0.29<br>(0.30) | 0.40<br>(0.41) |
| B) Males | i) $r_p$ | 0.40<br>(0.51) | 0.40<br>(0.51) | 0.42<br>(0.53) | 0.68<br>(0.72) | 0.43<br>(0.53) | 0.60<br>(0.67) |
| | ii) $r_s$ | 0.45<br>(0.54) | 0.52<br>(0.63) | 0.52<br>(0.64) | 0.70<br>(0.74) | 0.50<br>(0.62) | 0.70<br>(0.74) |
| | iii) B | 0.58<br>[1.31]<br>(0.48)<br>[1.10] | 0.14<br>[0.87]<br>(0.14)<br>[0.87] | 0.24<br>[0.88]<br>(0.24)<br>[0.88] | 0.91<br>[1.07]<br>(0.92)<br>[1.07] | 2.09<br>[1.04]<br>(2.09)<br>[1.04] | 2.38<br>[1.08]<br>(2.38)<br>[1.08] |
| | iv) adj $R^2$ | 0.15<br>(0.29) | 0.29<br>(0.30) | 0.31<br>(0.31) | 0.54<br>(0.55) | 0.31<br>(0.32) | 0.47<br>(0.47) |
